# Predicting and Controlling Collective Fate in Multicellular Systems

**DOI:** 10.64898/2025.12.11.693804

**Authors:** Tiam Heydari, Omar Bashth, Joelle Fernandes, Bhavya Sabbineni, Daniel Aguilar-Hidalgo, Jessica Chen, Nika Shakiba, Leah Edelstein-Keshet, Peter W. Zandstra

## Abstract

Collective behavior is a defining property of multicellular systems, where coordinated outcomes emerge from local cell–cell interactions. Yet the quantitative rules linking single-cell decision-making to tissue-scale organization remain poorly resolved. Here, we develop a quantitative framework that defines an order parameter predicting when initially disordered colonies undergo a transition to ordered fate alignment and when minimal, localized inputs can redirect their collective state. This analysis reveals a distinct control regime in which multicellular assemblies become susceptible to a single engineered “guide” cell. We validate these predictions by introducing guide cells that integrate into unperturbed colonies and redirect fate patterns within the theoretically defined control windows. Together, these results connect single-cell decision rules to emergent tissue-level organization and establish a generalizable biological control strategy in which a minority engineered subset can reliably redirect the developmental trajectory of a much larger multicellular population.

Summary figure:
Emergence, Scaling, and Control of Multicellular Collective Order.
(**A**) Cell-number–dependent collective order. Sparse colonies lack coordination and yield disordered fate distributions; dense colonies exhibit coordinated, ordered outcomes. (**B**) Collective order rises with colony compactness (∝ cell density) and collapses across colony radii R onto a single curve. (**C**) Once coordination emerges, a single “guide cell” can steer fate-analogous to a sheepdog guiding a flock. The herding efficacy exhibits a biphasic window versus cell number.

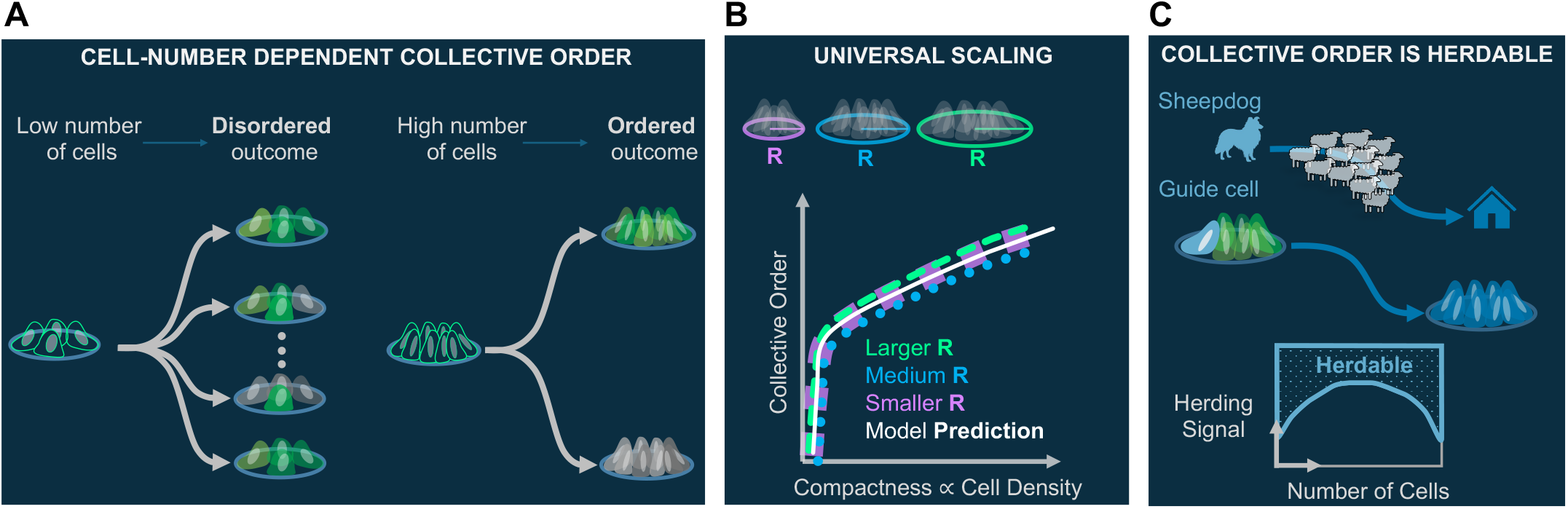

## Introduction

Collective behavior is a fundamental property of living systems, emerging when individual units interact through local rules to produce coordinated, large-scale outcomes. From bacterial swarms and immune cell coordination to the self-assembly of tissues during development, these phenomena arise from networks of interactions that catalytically propagate information across the collective (*1*). While the molecular logic of single-cell decision-making is increasingly well mapped (*2, 3*), the quantitative principles that translate individual cell states into ordered tissue-scale properties remain poorly understood. Without such principles, our ability to predict and control multicellular self-organization is limited.

Tissue formation arises from the integration of diverse information layers (*4*) - including gene regulatory programs (*5*), secreted morphogens (*6*), and mechanical forces (*7, 8*) - that together transform single cells into structured, functional assemblies (*9*). This multiscale integration drives the emergence of complexity and heterogeneity, with the outcome of orchestrated cell behavior depending as much on cell-cell interactions and environmental context as on intrinsic genetic cues (*10*). A central goal of regenerative medicine is to harness these principles to build functional tissues in a controlled manner, or to reprogram diseased tissue functions back to healthy states (*11*). The growing availability of single-cell and spatial atlases (*12, 13*) now offers detailed maps of the molecular states that define diverse tissues and developmental stages. However, these atlases are static snapshots; they tell us what is present but not how to translate those patterns into actionable “recipes” for guiding tissue-residing cells to the right states at the right time. Bridging this gap requires decoding the dynamical rules that connect molecular states, local interactions, and tissue-scale organization. Quantitative frameworks that link single-cell decision circuits, local communication, and tissue-level order would fill this longstanding gap, enabling the translation of descriptive cell atlases into predictive and programmable tissue control strategies.

Here we propose a foundational multiscale, data-driven framework that connects molecular decision circuits to collective tissue-level behavior. Our approach integrates bottom-up (*14*) and top-down (*15*) multicellular systems analysis, yielding a functional model that operates across scales. We implemented this framework in a controlled system using human pluripotent stem cells (hPSCs) confined to spatial patterns (*16*) and exposed to uniform morphogen stimulation. By combining imaging with time-resolved single-cell RNA-seq (scRNA-seq), we tracked how individual cells responded to a uniform cue, how they communicated with neighbors, and how fate decisions propagate across scales.

Single cell RNA-seq analysis revealed that, under uniform stimulation, hPSC fate bifurcated into two distinct transcriptional states. We formulated an order parameter (*17*) to quantify collective alignment and inferred a dense interaction network linking ligand-receptor pairs between fate-defined populations. From these data, we developed a minimal functional model that quantitatively captured the emergence of ordered fate patterns across colony sizes and geometries. The model predicted, and experiments confirmed, a biphasic susceptibility to external guidance. Colonies were more steerable either when very small, where a single cell could dominate, or large, where collective behavior has emerged. By engineering synthetic guide cells, we demonstrated that a single engineered cell could steer colony fate only within the predicted regimes, revealing a minimal, local strategy for controlling multicellular decision-making.

These results establish a generalizable framework for linking single-cell decisions, local communication, and tissue-scale organization, and demonstrate that multicellular fate decisions can be both emergent and controllable. This work provides a blueprint for programming multicellular systems through minimal, localized interventions, offering new strategies for synthetic biology (*18*), regenerative medicine (*19*), and adaptive tissue engineering (*20*).

## Results

### Time-resolved scRNA-seq reveals bifurcating responses to cytokine in confined stem cell-driven colonies

To investigate how cells coordinate fate decisions, we used hPSCs as a model system. These cells can recapitulate some aspects of the earliest stages of embryonic development, particularly where important fate decisions occur in the context of few neighboring cells (*21*). We seeded hPSCs onto light-activated adhesive patterns to create spatially confined colonies (*16, 22*) 35μm in radius. Colonies were uniformly stimulated with low-dose BMP4 (0.5 ng/ml), a developmental morphogen known to trigger cell state transition and differentiation (*23*) and imaged 42 h later using SOX2-GFP reporters, where SOX2 marks pluripotency at 0h (**Fig. 1A, S1**, and **S2**). Strikingly, despite identical initial conditions, individual colonies diverged to distinct outcomes: some retained SOX2 expression, others down-regulated it, and some exhibited a mixture of cell compositions (**Fig. 1B**).

**Fig. 1.**
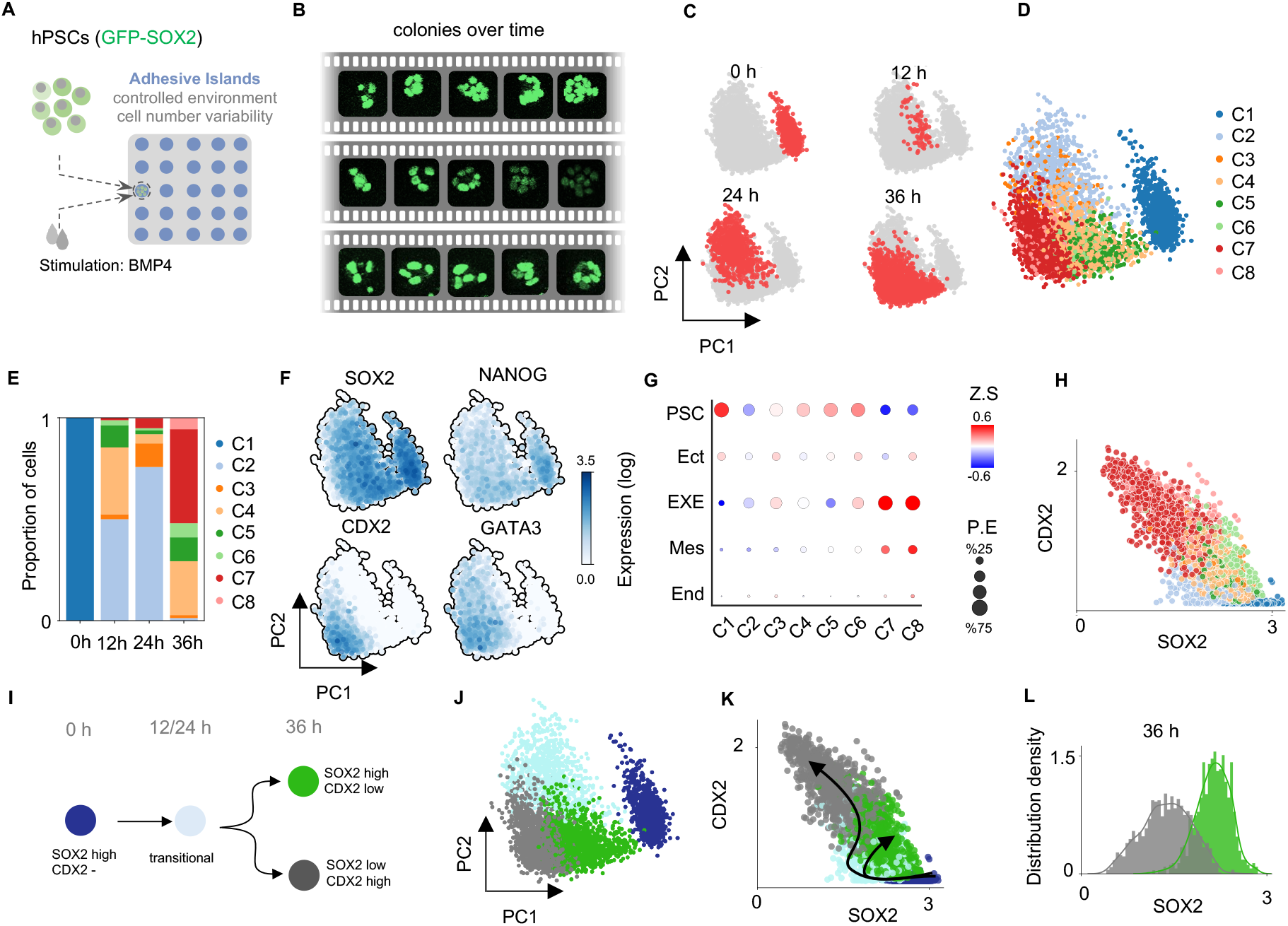
A confined stem cell system with uniform cytokine stimulation reveals a bifurcation-like transition to mutually exclusive fates. (**A**) Experimental setup: hPSCs are randomly seeded on micropatterned (MP) substrates and uniformly stimulated with low-dose BMP4 to induce differentiation in confined geometries. (**B**) Time-lapse imaging of SOX2-GFP expression in three representative colonies shows distinct population-level outcomes, including homogeneous maintenance, complete loss, or mixed SOX2 expression. (**C**) Principal component projection of single-cell transcriptomes at 0-, 12-, 24-, and 36-h post-stimulation reveals continuous progression over time. (**D**) Unsupervised clustering identifies eight distinct transcriptional states across the time course. (**E**) Bar plot showing the proportion of cells assigned to each cluster at each time point. (**F**) Expression of selected marker genes (SOX2, NANOG, CDX2, GATA3) projected onto the PC space illustrates mutually exclusive fate signatures. (**G**) Dot plot summarizing lineage program resemblance based on curated marker genes across clusters, dot size reflects the percentage of cells expressing markers (P.E), and color denotes the ZScroe (Z.S), (see Supplementary Methods for detail). (**H**) Scatter plot of SOX2 versus CDX2 expression, colored by cluster, shows that this two-gene axis alone robustly resolves major cell states (log transformed). (**I**) Schematic of fate progression: cells begin in a SOX2-high/CDX2-negative state, pass through a transitional population, and bifurcate by 36 h into SOX2-high/CDX2-low or SOX2-low/CDX2-high terminal states. (**J**) PC space colored by expression-defined fate groups; colors represent cell states in (I). (**K**) Same grouping visualized in SOX2–CDX2 space, with arrows indicating progression of cell states (log transformed). (**L**) Density plot of SOX2 mRNA expression in the two terminal populations at 36 h (SOX2-high/CDX2-low in green and SOX2-low/CDX2-high in grey) shows minimal overlap, supporting the use of SOX2-GFP as a proxy for final fate identity.

To determine whether the observed SOX2 dynamics reflected *bona fide* cell state transitions rather than transient gene expression fluctuations, we performed time-resolved single-cell RNA sequencing on geometrically confined colonies under uniform stimulation. Cells were sampled at 0, 12, 24, and 36 h post-stimulation to capture the temporal progression of transcriptional states. Dimensionality reduction of transcriptomic data revealed progression in principal component (PC) space over the course of 36 h (**Fig. 1C**). Unsupervised clustering identified eight transcriptionally distinct states (**Fig. 1D**). C1 was enriched for cells at 0 h, and C4-C8 for 36 h, corresponding to terminal outcomes. Analysis of lineage marker expression (*24*) showed that C1 expressed pluripotency genes including SOX2 and NANOG, while C2 and C3 displayed an intermediate phenotype with partial downregulation of pluripotency markers and emerging extraembryonic signatures. C4-C6 re-expressed pluripotency markers with minimal lineage activation, suggesting the maintenance of PSC-like state. C7 and C8 upregulated extraembryonic genes, including CDX2 and GATA3, indicating a shift toward extraembryonic-like identity (**Fig. 1F, G**, and **S3**).

The SOX2-CDX2 expression axis alone robustly separated pluripotent and differentiated states (**Fig. 1H**). Grouping clusters along this axis revealed a shared cell state transition trajectory where cells began in a SOX2-high/CDX2-negative state at 0 h, progressed through a transitional population at 12–24 h, and by 36 h bifurcated into either a SOX2-high/CDX2-low or SOX2-low/CDX2-high fate (**Fig. 1I-K**). Notably, at 36 h, SOX2 expression alone was sufficient to distinguish the two terminal populations with minimal overlap, despite potential dropout in single-cell transcriptomic data (**Fig. 1L**). These results validate SOX2-GFP fluorescence as a reliable reporter of fate identity in this system, enabling tracking of multicellular fate dynamics and setting the stage for analysis of potential higher order (collective) decision-making.

### Emergence of collective order through cell-cell communication

Under uniform cytokine stimulation, confined hPSC colonies bifurcated into distinct fates. The origin of this variability was unclear: did it arise from cell-intrinsic stochasticity (*25*) within colonies, or from coordinated fate decisions at the colony level? To distinguish between these possibilities, we first confirmed that neighboring colonies act independently (**Fig. S4**, and **S5**), ruling out long range spatial cues. We then defined an Order Parameter (𝒪_p_) to quantify the degree of fate alignment within each colony. In this framework, colonies with fully random outcomes among cells show low order (𝒪_p_ ≈0) while those with coordinated and consistent outcomes show high order (𝒪_p_ ≈1), (**Fig. 2A** and **S6**). Applying this metric across colonies with varying numbers of cells showed that 𝒪_p_ increased as the number of cells per colony grew (**Fig. 2B** and **S7**). Thus, rather than increasing randomness, higher cell numbers were associated with a transition from less ordered, potentially more stochastic behavior to more coordinated collective outcomes.

**Fig. 2.**
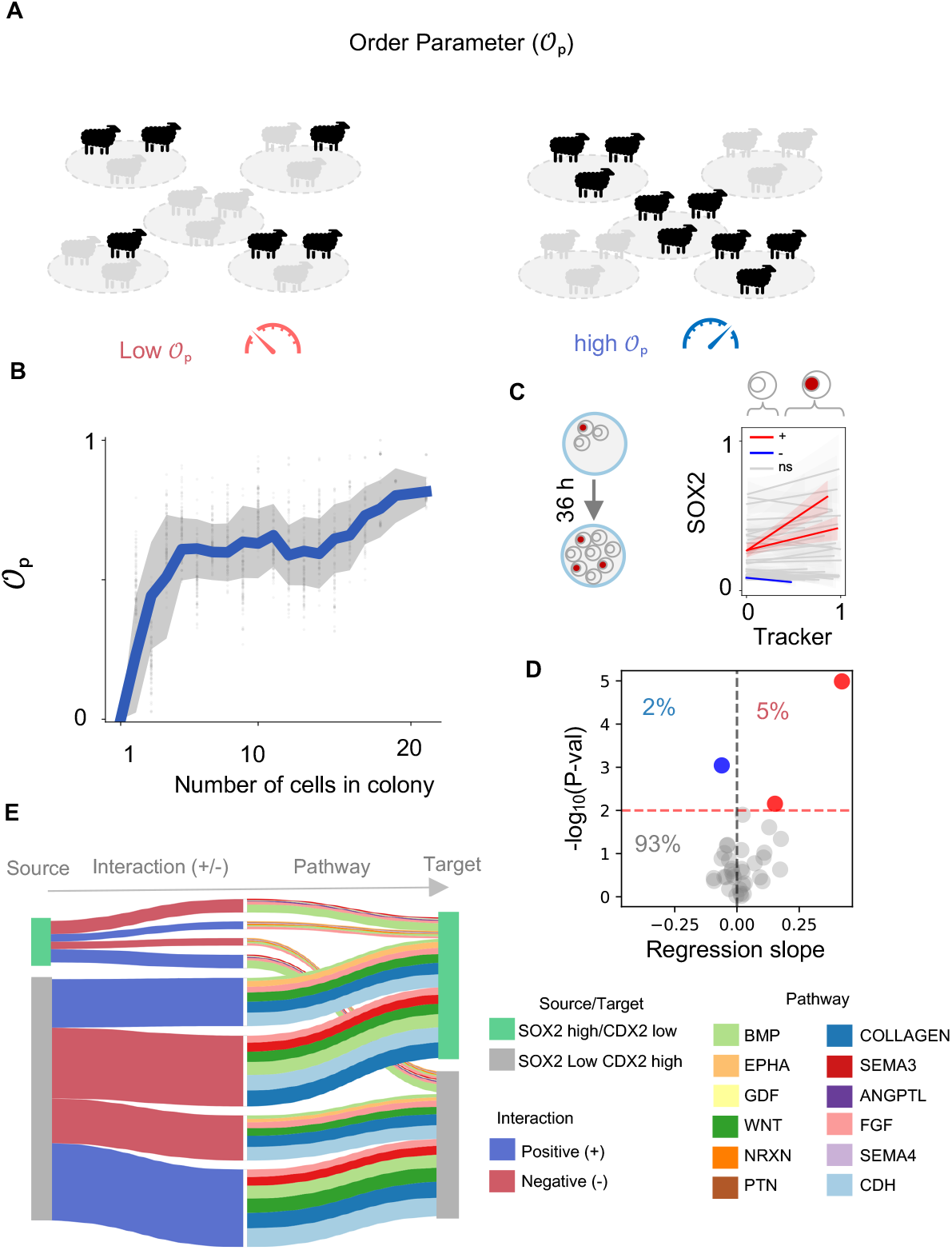
Collective fate decisions emerge from intercellular signaling. (**A**) Conceptual illustration of the Order Parameter (𝒪_p_), used to quantify intra-colony coordination. Colonies in which all cells adopt the same fate exhibit high order, while those with mixed fates show low order. (**B**) Order increases with the number of cells per colony, indicating that collective alignment strengthens with group size. Shaded region shows mean ± s.d. across biological replicates. (**C**) Left: Schematic of lineage-tracing experiment using labeling. Right: SOX2 intensity at 36 h versus cell label intensity across individual cells within colonies. Lack of strong correlation suggests that fate is not clonally inherited. (**D**) Summarizing the regression analysis from (C). Each dot represents a colony, showing regression slope versus statistical significance. Most colonies show no significant correlation, ruling out clonal dominance as a general mechanism of coordination. (**E**) Inferred ligand– receptor signaling network based on single-cell transcriptomic data. Putative communication routes are grouped by the source and target population, including self-targeting interactions. The network reveals extensive reciprocal signaling between SOX2-high/CDX2-low and SOX2-low/CDX2-high populations, involving both activating (blue) and inhibitory (red) interactions. Even after filtering, many candidate pathways remain (shown by colors in the right half of the plot), highlighting the complexity of intercellular communication and raising the question of which signaling modes may underlie the coordinated fate decisions in more crowded colonies.

One possible explanation for the coordinated fate outcomes observed within colonies is clonal dominance, where descendants of a competent single cell adopt the same fate due to inherited internal states (*26*). To test this, we labeled ∼5% of cells at the beginning of the experiment with a fluorescent tracker and looked at their progeny over 36 h as they divided (**Fig. 2C**). Most colonies showed no association between tracker signal and SOX2 levels, and statistical analysis revealed that most colonies had no significant correlation (**Fig 2D**). Only a small minority showed positive or negative trends. These findings discount clonal inheritance as the dominant driver of fate coordination, instead implicating dynamic intercellular mechanisms such as signaling-mediated communication.

To investigate intra-colony signaling mechanisms, we constructed ligand-receptor interaction networks from the scRNA-seq data, stratified by cell identity (*27*). This analysis revealed dense networks of potential signaling interactions, including BMP and FGF ligands produced by one fate population and cognate receptors enriched in the other (**Fig. 2E** and **S8**). The prevalence of reciprocal signaling motifs suggested the presence of positive (and negative) feedback loops capable of reinforcing (or attenuating) shared fate identity (*28*). Together, these results indicate that the transition from low to high order with increasing colony size likely arises from local communication and feedback, enabling colonies to behave as coordinated collectives rather than independent cells (*28*).

### A minimal model of inter- and intra-cellular dynamics

To bridge molecular and cellular information across scales, we first examined the internal state dynamics of individual cells over the course of the experiment using scRNA-seq. At the beginning, cells were uniformly in a SOX2-high/CDX2-state. By 36 h, the population had diverged into two high-density regions in SOX2-CDX2 space, corresponding to either a SOX2-high/CDX2-low or SOX2-low/CDX2-high state (**Fig. 3A**). This distribution suggests the presence of a bi-stable switch governing fate commitment. To mimic this behavior, we constructed a minimal model of a single cell with two mutually inhibitory states (*29*), (**Fig. S9a**). Starting from a SOX2-high/CDX2-state, the system evolves toward one of two stable states over time when destabilized. Either maintaining SOX2 and suppressing CDX2, or vice versa. This minimal toggle-switch model recapitulated the experimentally observed bifurcation in expression space (**Fig. 3B**), providing a simple framework to study the dynamics of fate resolution at the single-cell level.

**Fig. 3.**
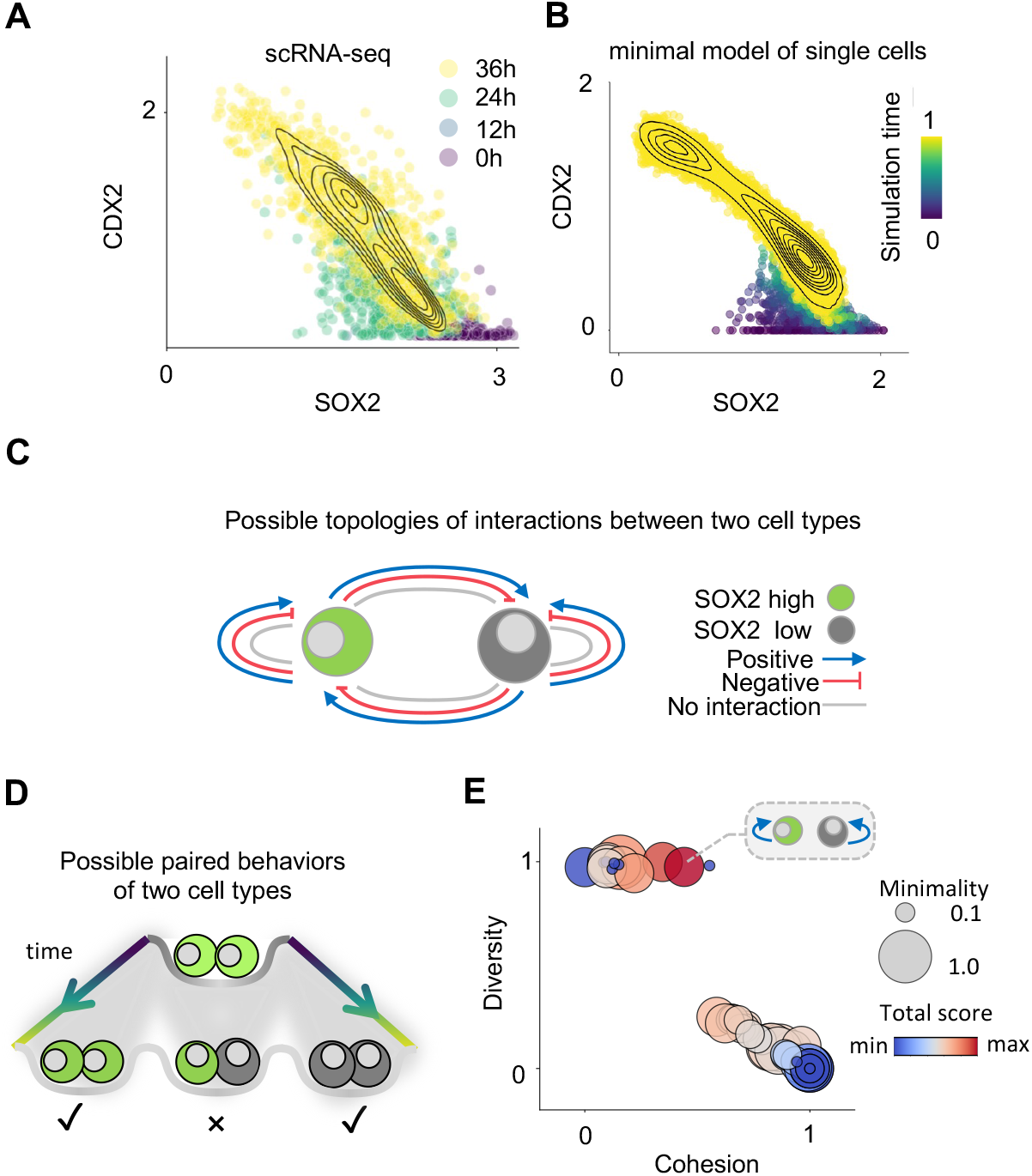
A minimal model to capture cellular dynamics. (**A**) Single-cell RNA-sequencing of cells over 36 h, projected into SOX2–CDX2 expression space and color-coded by time. By the end of the experiment, cells bifurcate into two distinct fates: SOX2-high/CDX2-low and SOX2-low/CDX2-high, as shown by kernel density contours (log transformed). (**B**) Minimal model simulation reproduces the bifurcating trajectories seen in (A), with simulated cells transitioning from a common starting point to one of two stable states. Contours mark final state distributions, confirming the model’s ability to recapitulate fate divergence. (**C**) Schematic of all possible pairwise signaling topologies between SOX2-high (green) and SOX2-low (gray) cells. Each directed edge encodes positive (blue), negative (red), or absent (gray) interactions, yielding 81 possible configurations. (**D**) Schematic illustration of possible paired behaviors for two cells over time. The system begins with identical starting conditions and diverges into distinct fates. Only symmetric outcomes-where both cells adopt the same terminal state (green-green or gray-gray) are considered coordinated and fate-aligned (✓). Mixed outcomes (green-gray) are disordered and fail to reflect collective alignment (✗). The dominant signaling topology must reliably produce these fate-coordinated outcomes under uniform stimulation. (**E**) Systematic evaluation of all topologies across three axes: diversity (y-axis, capacity to support both fates), cohesion (x-axis, degree of alignment within colonies), and minimality (circle size, inverse of interaction count). Color reflects overall score (red = high, blue = low). A small subset of topologies emerges as both simple and functionally robust.

To connect the intracellular dynamics with intercellular signaling, we considered the possible modes of communication between the two cell states. As shown in **Fig. 3C**, each state can signal to itself or to the other state through either positive, negative, or neutral interactions, yielding a total of 3^4^ = 81 possible signaling configurations. While multiple modes co-exist with varying strengths, we asked whether a dominant signaling topology exists in our system as the driver of the observed coordinated fate alignment (**Fig. 2B**). This implies that the relevant signaling mode must produce cohesive and collective behavior across colonies in both fate branches (**Fig 3D**). We systematically simulated all topologies and evaluated each based on three criteria: diversity (ability to support both fates across conditions), cohesion (internal fate alignment within colonies), and minimality (number of signaling links required). This allowed us to identify a high-scoring model that produced robust coordination and fate divergence using simple interaction rules (**Fig 3E** and **S10**).

### A functional model predicts collective fate coordination

While our previous analysis focused on the topology of signaling between pairs of cells, colonies consist of many cells communicating in a confined, shared space (*30, 31*). To scale up the model, we connected multiple cells using a geometric dilution that links the number of signal-producing cells to the volume available (**Fig. S11**). This accounts for how signaling density increases with increasing cell number under circular pattern confinement. The full model includes a single free parameter controlling the strength of intercellular signaling, which we fit using data from colonies on 35 μm radius (R) adhesive islands (**Fig. S12** and **S13**). The fitted model accurately reproduced the experimental relationship between colony size and 𝒪_p_, showing that 𝒪_p_ increases with the number of cells (**Fig. 4A**). This behavior reflects a threshold effect: once a colony reaches a critical number of cells, cumulative signaling overrides intrinsic variability and enables a collective fate switch. To test whether fate coordination depends on intact intercellular signaling, we inhibited the BMP pathway in the middle of the experiment via a BMP inhibitor (BMP is the mediator of the initial differentiation stimulus and endogenous communication, see Supplementary Methods). In R=35 μm colonies, BMP inhibition disrupted fate alignment and reduced the order parameter, consistent with model predictions **(Fig. 4B**).

**Fig. 4.**
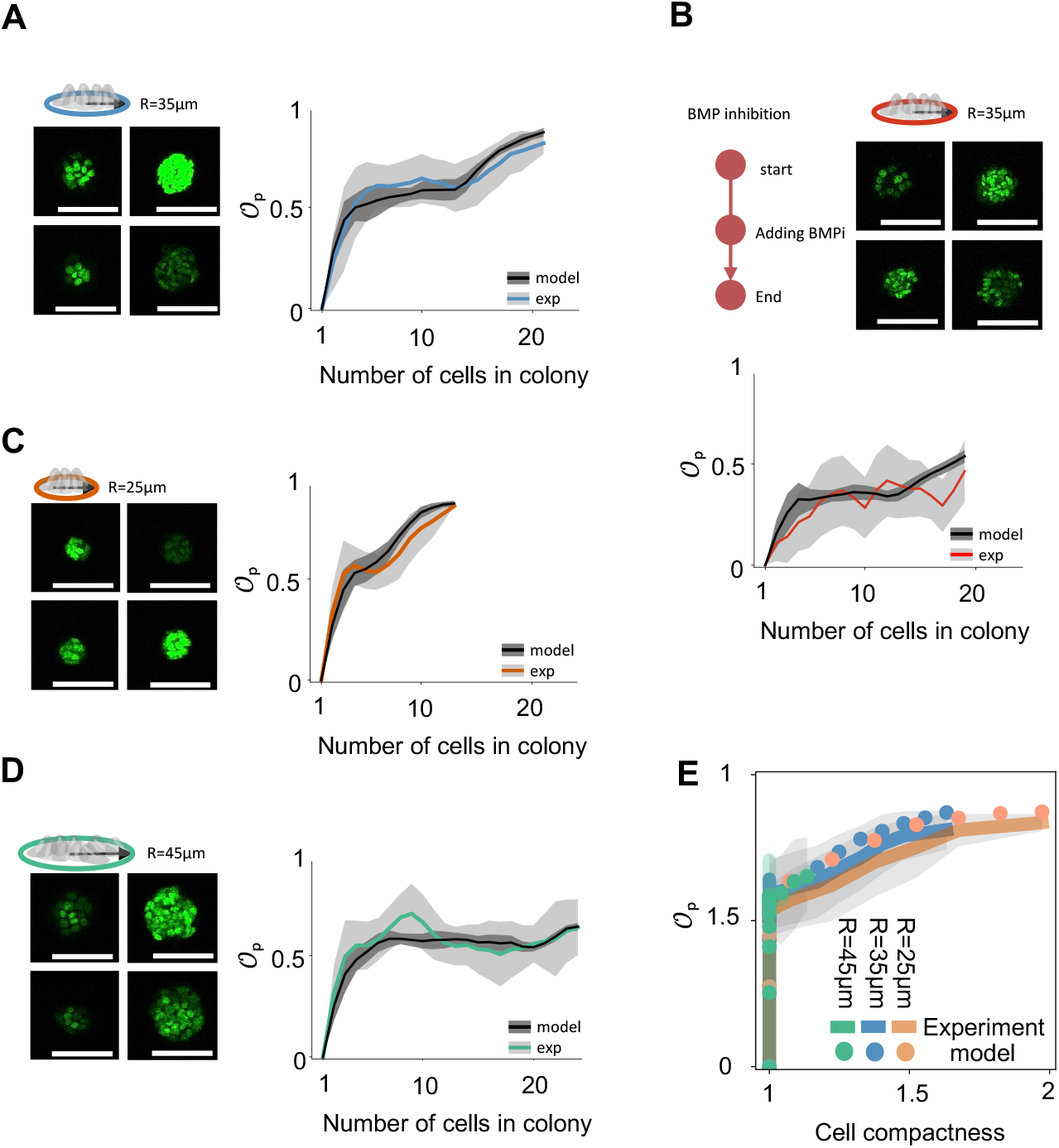
A functional model predicts collective cell behavior across conditions. (**A, C, D**) Experimental validation of model predictions across three colony sizes: R = 35 μm (used in model fitting), R = 25 μm (g), and R = 45 μm (h). Left: representative confocal images of SOX2^+^ (bright) and SOX2^−^ (darker) cells. Right: plots of the order parameter (𝒪_p_) versus cell number. Experimental data (colored lines ± s.d.) and model predictions (black line) show strong agreement. Notably, coordination emerges at lower cell numbers in smaller colonies, while in larger colonies (R = 45 μm), coordination remains partial even at high cell counts. Scale bars show 100μm. (**B**) Disruption of intercellular signaling using a BMP pathway inhibitor (BMPi) in R = 35 μm colonies. Left: SOX2 expression patterns show disrupted fate alignment. Right: BMPi-treated colonies (red) show reduced order (𝒪_p_), matching model predictions under weakened signaling and confirming the role of feedback in coordinating fate. Scale bars show 100μm. (**E**) Universal scaling with compactness. Plotting 𝒪_p_ against cellular compactness (cell number per colony area) reveals a single unifying trajectory across all colony sizes. Experimental data and model predictions collapse onto the same curve, suggesting compactness governs the onset of multicellular order irrespective of absolute colony size.

To test the model’s generality, we asked whether it could predict fate coordination across colonies of different sizes without further parameter tuning. Because colony diameter influences both the number of interacting neighbors and the extent of spatial confinement, changes in geometry are expected to impact collective behavior. Remarkably, the same model successfully predicted the emergence and saturation of order in both smaller (R = 25 μm) and larger (R = 45 μm) colonies (**Fi. 4C** and **D**). In smaller colonies (R=25 μm), coordinated fate alignment emerged at lower cell numbers, consistent with stronger effective signaling due to tighter spatial packing and reduced diffusion volume. Conversely, in larger colonies (R=45 μm), even with many cells, the model correctly predicted that coordination remained partial and never reached the levels observed in geometrically smaller colonies (**Fig. S7**). These results highlight that cell intrinsic networks, extrinsic signaling interactions, and biophysical constraints are coordinated across scales to collectively specify fate outcomes in multicellular systems.

We next asked whether cellular compactness alone (defined to be proportional to the cell density, see Supplementary Methods) could explain the emergence of collective order across all tested conditions. We plotted the order parameter (𝒪_p_) against compactness; both model predictions and experimental measurements from all three colony sizes collapsed onto a single trajectory (**Fig. 4E**). This universal scaling suggests that compactness acts as a unifying variable, capturing the interplay between cell number and spatial constraint that governs multicellular coordination, independent of absolute colony size, or number of cells in this context.

### Harnessing collective behavior to steer fate decisions

Having established density-dependent collective behavior, we next asked how information originating from a single cell propagates through the group. To address this, we turned to our model (**Fig. 5A**) and defined an induction factor, χ, which quantifies how the signaling state of a colony changes in response to the state of a single cell (**Fig. S14**). Simulations revealed a biphasic pattern: χ was low in intermediate-sized colonies but increased in both small and large colonies (**Fig. 5B**). A bi-sigmoid fit to the data highlighted two transition points, marking the boundaries where responsiveness switched regimes. As expected, in small groups, a single cell has significant influence by constituting a large fraction of the population. Unexpectedly, however, in large groups, coordinated multi-scale dynamics propagate the signal. In intermediate groups, induction is diluted.

**Fig. 5.**
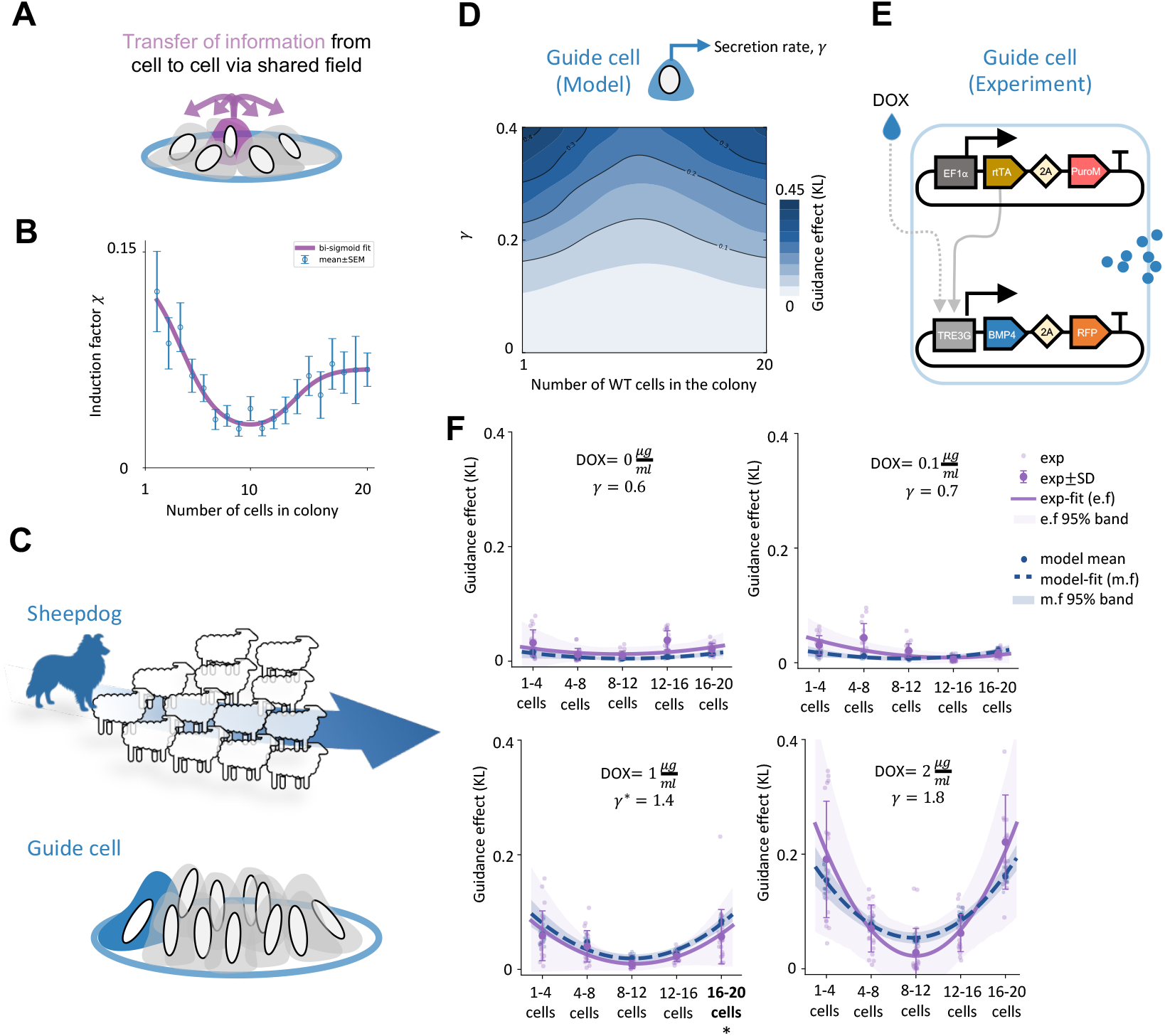
A single guide cell can steer collective fate decisions. (**A**) Schematics of cells influencing each other via a shared signaling field. (**B**) Model prediction of colony-level induction factor, χ as a function of colony size. χ measures the sensitivity of colony signaling state to individual cell. Simulations predict biphasic responsiveness: low χ at intermediate sizes but high χ for very small or large colonies. purple line shows a bi-sigmoid fit to the data. (**C**) Drawing conceptual analogy. A single agent (e.g., a shepherd dog) can control a large group once collective motion emerges (top). We hypothesize that an engineered “guide cell” (blue) might similarly direct fate in a dense colony of cells (bottom) when collective behavior emerges. (**D**) Phase plot of guide cell model predictions for a BMP-secreting guide cell that emits signal at rate γ. Guidance effect (KL divergence between SOX2 distributions with vs without one guide cell) is strongest at sufficiently high γ in the smallest and largest colonies, and weakest at intermediate sizes. (**E**) Experimental design. Guide cells were engineered via lentivirus transduction to carry a DOX-inducible TRE3G–BMP4 cassette with an RFP reporter. (**F**) Experimental guidance measurements across DOX doses (0, 0.1, 1, 2 μg mL^−1^). Colonies containing exactly one guide cell are compared to matched colonies without a guide cell. A biphasic guidance pattern emerges at higher DOX (1–2 μg mL^−1^). Calibrating guide cell model signal section γ using the 16-20 cell group at DOX=1 μg mL^−1^ allows the model to predict all other size-dose conditions.

Inspired by parallels in nature, we turned to an analogy from collective animal behavior (*32–34*). Just as a single sheepdog can steer a flock once collective motion emerges (*35*), we hypothesized that an engineered “guide cell” could direct (*36*) the gene-expression fate of neighboring cells in dense colonies (**Fig. 5C**). We first evaluated this idea by modeling the effect of a guide cell that secretes BMP at the rate of γ. We quantified the effect of guidance as the Kullback-Leibler divergence (*37, 38*) between SOX2 distributions in colonies with and without a guide cell (see Supplementary Methods and **Fig. S15**). Model predictions recapitulated the biphasic signature: when the secretion rate is above a threshold, guidance starts to appear in small and large colonies, with less effect at intermediate sizes (**Fig. 5D**).

To test this prediction experimentally, we engineered synthetic guide cells by placing BMP4 under the control of a doxycycline (DOX)-inducible promoter (*39*), coupled with a reporter (**Fig. 5E** and **S16**). We introduced a small proportion of guide cells (15%) into colonies of WT cells and selected colonies with either one guide cell or no guide cell for downstream analysis. Guidance effects were most significant at the higher DOX dose (1-2 μg/mL), and more distinguishably in the smallest (1-4 cells) and largest (16-20 cells) colonies (**Fig. 5F**). By fitting signal secretion γ to 16-20 cells colonies at DOX=1μg/mL (**Fig. S17**), the model predicted all other conditions (cell numbers and concentrations), (**Fig. 5F**).

Together, these results show that a single engineered cell can steer collective fate decisions within predictable control windows. This reveals a general principle: multicellular systems can be programmed through minimal strategically placed inputs, transforming collective behavior into a controllable design feature.

## Discussion

Altogether, our results show how multicellular fate decisions emerge from local interactions and can be steered with minimal interventions. Our model bridges molecular-scale gene expression variability to multicellular behavior within a controllable platform. Guided by scRNA-seq, we reduced cells’ dynamics to a tractable mathematical framework. This abstraction is context-dependent and omits additional possible fate axes of hPSCs (*40*); therefore, its parameters should not be interpreted as universal.

To recover ordered states, we performed a systematic top-down search across possible interactions, which led us to identify a minimal set of rules. This framework parallels classical models of self-organization, including the Vicsek model of collective migration (*41*), spin alignment using the Ising model (*42*), and cellular automata (*43*) where simple local rules give rise to emergent order. Importantly, our model combines the complex multiscale aspects of cellular interactions, biochemical, mechanical, and electrical signaling, into a representation that captures the key determinants of collective behavior while remaining experimentally testable (*44, 45*).

Future studies should extend our framework to three-dimensional tissue architectures, where adhesion, migration, and diffusion may additionally influence collective fate decisions. A central question in developmental biology remains how simple local rules give rise to the organized structure of tissues and organs. Our framework could be applied to developmental systems such as embryos (*46*) or organoid platforms (*47*) particularly when integrated with high-throughput, time-resolved spatial transcriptomic data. Such applications may require new modeling and analytical approaches that capture additional layers of biophysical and signaling complexity.

Our model revealed effective information transfer in the collective phase, and both theory and experiment showed that a single “guide cell” can steer the collective dynamics. This finding echoes natural leadership paradigms: in collective animal migration, small groups of informed individuals can direct the motion of the whole (*36*); in epithelial wound healing, leader cells at the front coordinate migration and are later eliminated by cell competition once closure is achieved (*48*); and in early embryogenesis, the Spemann organizer directs germ-layer specification through coordinated migration and signaling (*49*). Extending this framework to three-dimensional tissues could enable the design of engineered guide cells that respond to defined stimuli (*50*), can be positioned within multicellular assemblies, and eliminated once patterning is complete. Such an approach would expand the experimental and therapeutic toolkit for directing tissue organization and regeneration.

In conclusion, we define experimentally testable principles for how collective multicellular fate decisions arise and can be steered. By quantitatively linking single-cell stochasticity to mesoscale communication and colony-level order, the work provides predictive rules for multicellular organization. The framework connects developmental decision-making to broader classes of collective systems while acknowledging simplifying assumptions such as confined geometries, a restricted signaling axis, and coarse-grained dynamics. These constraints clarify where predictions are expected to hold and motivate direct tests in systems with more complex architectures. More broadly, the framework provides a basis for comparative analyses across developmental and engineered tissues and for identifying minimal interventions capable of reshaping collective tissue outcomes.

## Acknowledgments

The authors thank C. Zimmerman for assistance with cell culture and routine quality control, T. Stach for help with single-cell RNA sequencing, and R. D. Jones for support with sequencing sample preparation and for critically reading the manuscript. We also gratefully acknowledge the assistance and support of our laboratory colleagues and collaborators.

## Author contributions

T.H. and P.W.Z. conceptualized and designed the study. T.H., D.A.H., and P.W.Z. wrote the first draft; T.H, P.W.Z, L.E.K, and N.S. reviewed and edited the manuscript. T.H., J.F., O.B., and B.S. performed cell culture and experiments on adhesive islands. T.H. and J.F. developed and performed biomanufacturing of adhesive islands. T.H. and B.S. performed single-cell sequencing. T.H., J.F., and O.B. performed imaging. T.H. developed mathematical models, conducted in-silico simulations, and performed bioinformatics analyses. T.H, O.B., and N.S. designed the synthetic guide-cell experiments, and O.B. designed and generated the DOX-inducible synthetic guide-cell line. J.C and L.E.K designed the Morpheus simulations. T.H., J.C., and D.A.H. conducted analyses of simulated data. P.W.Z supervised the study.

## Data and materials availability

Data supporting the findings of this study are available in the main text or the supplementary materials. All single-cell RNA-seq data and associated code will be deposited in public repositories and made available upon publication.

## Supplementary Materials

Materials and Methods

Supplementary Text

Figs. S1 to S17.

References (51-79)

## Supplementary Materials

### 1. Materials and Methods: Experiments

#### 1.1 Routine cell culture and quality control

##### Routine cell culture

AICS-0074-026, a specific human induced pluripotent stem cell (iPSC) line called WTC-mEGFP-SOX2-cl26 (*51*) was used for experiments. Cells were maintained in 6-well plates coated with Geltrex (1:50, Thermo Fisher) and cultured in mTeSR1 medium (STEMCELL Technologies), with daily medium replacement. Cultures were passaged every 3-4 days at ∼75-80% confluency using StemPro Accutase (Thermo Fisher) following the manufacturer’s instructions, typically at a 1:12-1:18 split ratio. ROCK inhibitor (Y-27632, 10 µM; STEMCELL Technologies) was added for 24 h after passaging to enhance and promote survival. Cells were regularly tested for mycoplasma contamination and morphological quality control (**Fig. S1 a-c**).

##### Quality control

iPSCs were harvested using TrypLE (Thermo Fisher #12605208) to obtain a single-cell suspension, followed by centrifugation at 1000 rpm for 5 min and washing in Hank’s Buffered Salt Solution containing 2% FBS (HF), (Thermo Fisher #14175103). Approximately 2 × 10^5^ cells were resuspended in 100 µL HF per sample, including unstained and secondary-only controls. Cells were fixed by adding an equal volume of 8% paraformaldehyde (PFA) to achieve a final concentration of 4% and incubating for 15 min at room temperature with vortexing. Fixed cells were washed, resuspended in HF, and permeabilized with 100 µL ice-cold 100% methanol for 2 min. Following a wash, cells were incubated in 100 µL HF containing mouse anti–OCT-3/4 (BD Biosciences #611203; 1:200 dilution) for 20 min at room temperature. After washing, secondary staining was performed using donkey anti-mouse IgG AlexaFluor 647 (Thermo Fisher #A31571; 1:200 dilution) for 20 min at room temperature in the dark. Rabbit anti-Sox2 staining was omitted due to endogenous SOX2-EGFP fluorescence. Cells were washed twice with HF and resuspended in 200 µL HF for flow cytometry analysis. Routine QC ensure 95%+ of cell as double positive (**Fig. S1 d**).

#### 1.2 Preparation of adhesive islands

Patterned substrates were prepared on 22 × 22 mm glass coverslips (VWR) placed in 35 mm tissue culture dishes. To minimize non-specific binding, coverslips were passivated with 5 mg/mL Lipidure-CM5206 (NOF America) in 100% ethanol. Circular adhesive islands were generated using custom quartz photomasks (Photo Sciences Inc.). Photomasks were cleaned in a plasma cleaner (Herrick Plasma; 3 minutes at 0.4 mTorr), and patterned coverslips were transferred onto coverslips by photo-oxidation with a UV-ozone cleaner (Jelight, for 30 min). After excessive washing with ddH_2_O, patterned regions were activated for ECM protein immobilization by first soaking in ddH_2_O for 30 min, then incubating with N-(3-Dimethylaminopropyl)-N′-ethylcarbodiimide hydrochloride (EDC; Sigma) and N-hydroxysuccinimide (NHS; Sigma) for 20 minutes.

Coverslips were rinsed twice with ddH_2_O and incubated overnight at 4 °C with Laminin-521 (STEMCELL Technologies, 5-10 µg/mL). Before cell seeding, Laminin-521 was removed by six washes with ice-cold DPBS containing calcium and magnesium (DPBS++, Fisher Scientific).

#### 1.3 Seeding, differentiation, and cell tracker

For experimental seeding, iPSCs were dissociated with Accutase (37 °C, 5-7 minutes) to generate single-cell suspensions, centrifuged, and resuspended in mTeSR1 supplemented with 10 µM ROCK inhibitor (Y-27632) and 1% penicillin/streptomycin (Invitrogen). Cells were plated at ∼50,000 cells per well to promote the formation of well-defined colonies (**Fig. S2**). After 5-6 h, the medium was replaced with fresh mTeSR1 containing penicillin/streptomycin but without ROCK inhibitor and with the indicated cytokine for differentiation (typically BMP4 at 0.5 ng/mL). Colonies were cultured for up to 42 h, at which point they were fixed for imaging with 4% paraformaldehyde (Thermo Fisher) in PBS for 15 min at room temperature, followed by three PBS washes. For clonal lineage analysis, ∼5% of cells were pre-labeled with CellTracker dye (Thermo Fisher) before seeding. Briefly, dissociated cells were resuspended in serum-free medium containing 1-5 µM CellTracker dye, incubated for 20-30 min at 37 °C, then washed twice in fresh medium to remove excess dye. Labeled cells were mixed with unlabeled cells at the indicated proportion and seeded as above.

#### 1.4 Single cell RNA sequencing

To profile transcriptional dynamics under spatial confinement, hPSCs were cultured on R=35 µm patterned substrates and stimulated with BMP4 (0.5 ng/mL). Cells were harvested at 0, 12, 24, and 36 h post-stimulation. At each time point, four patterned chips were dissociated with Accutase (37 °C, 9 min), quenched with mTeSR1 + penicillin/streptomycin, and pelleted (500 × g, 10 min). Cell viability was assessed using AOPI staining and an automated cell counter to ensure above 90% viability at this stage. Pellets were fixed overnight at 4 °C in 10x Genomics Fix Buffer (100 µL Fix Buffer, 125 µL 32% paraformaldehyde, 775 µL RNase-free water per mL). After 24 h, cells were centrifuged at 300 × g for 5 minutes, resuspended in 10x Genomics Quench Buffer (125 µL 8× Quench Buffer + 875 µL chilled RNase-free water), supplemented with 100 µL 10x Genomics Enhancer (pre-warmed to 60 °C), and adjusted to 10% glycerol before storage at -80 °C. Libraries were prepared using the Chromium Fixed RNA Profiling kit (10x Genomics) with the Human Probe Set according to the manufacturer’s instructions. Libraries were sequenced on an Illumina NextSeq 2000. Raw sequencing data were processed with Cell Ranger (10x Genomics, v7.1.0; Martian v4.0.10) using the cellranger count pipeline with the FRP human probe set reference (GRCh38). The resulting filtered gene-cell count matrices were used for downstream analysis.

#### 1.5 BMP pathway inhibition

To disrupt endogenous BMP signaling, colonies were treated with the selective BMP type I receptor inhibitor LDN-193189 hydrochloride (STEMCELL Technologies, Cat# 72147). LDN was added at 20 h post-BMP4 stimulation to a final concentration of 100 nM. Cultures were maintained under these conditions until the experimental endpoint at 42 h.

#### 1.6 Guide-cell experiment

Lentiviral constructs were designed to enable inducible BMP4 secretion. In construct vOB_9, the reverse tetracycline transactivator (rtTA) followed by T2A-puromycin was expressed under the EF1α promoter. In construct vOB_10v2, the BMP4 coding sequence (PCR-amplified from pCL050) was placed downstream of an inducible TRE3G promoter, followed by T2A-RFP670 as a fluorescent reporter. A blasticidin resistance cassette under the PGK promoter was included for selection. Both constructs were assembled in lentiviral backbones using Gibson Assembly (**Fig. S16a**).

Lentiviral particles were produced in Lenti-X 293T cells (Takara) cultured in DMEM supplemented with 10% FBS and 1% Glutamax. Approximately 4.5 × 10_6_ cells were seeded in 10-cm dishes one day before transfection. For each transfection, 1.64 pmol of either vOB_9 or vOB_10v2 was combined with 1.3 pmol psPAX2 and 0.72 pmol pMD2.G in 500 µL Opti-MEM, mixed with PEI (1:3 ratio), incubated for 15-20 min, and added dropwise to the cultures. The medium was changed the next day, and viral supernatants were harvested after 48 h, centrifuged, filtered through 0.45 µm filters, and stored at -80 °C. Viral titer for vOB_10v2 was estimated by transducing hPSCs with serial dilutions of viral supernatant in the presence of polybrene (8 µg/mL) and ROCK inhibitor (1 µM), followed by blasticidin selection (5 µg/mL). Cell counts after selection were used to construct a titration curve. For guide cells, hPSCs were first transduced with vOB_9 and selected with 0.5 µg/mL puromycin for three rounds to ensure rtTA expression. These cells were then transduced with vOB_10v2 at MOI 1, followed by three rounds of blasticidin selection (5 µg/mL). To assess inducibility, sender cells were treated with doxycycline (DOX; 0.01-10 µg/mL) for 24-48 h, and induction was monitored by RFP670 expression using fluorescence microscopy (ECHO Revolve) and flow cytometry (CytoFLEX), (**Fig. S16 b, c**).

Double-positive GFP^+^/RFP670^+^ cells were bulk-sorted on an Astrios cell sorter for use in micropattern experiments. BMP4 secretion was quantified by ELISA (Human BMP4 ELISA kit) following DOX induction. Sender cells were seeded in a 6-well plates, treated with DOX (0.0-3.0 µg/mL) for 48 h, and conditioned medium was collected. ELISA was performed according to the manufacturer’s protocol, with absorbance measured at 450 nm (**Fig. S16d**). Receiver cells were generated by transducing hPSCs with a TagBFP construct (**Fig. S16a**) and performing two rounds of FACS to ensure stable expression. Receiver-only controls were prepared in parallel.

For micropattern experiments, guide cells (MOI 1, FACS-purified) and receiver cells were propagated in mTeSR1 on Geltrex prior to use. On the day of seeding, cells were harvested with TrypLE, and were plated per micropattern slide at a 15:85 guide cell:receiver ratio in mTeSR1 containing 10 µM ROCK inhibitor. After 5-6 h, DOX was added together with a low BMP4 dose (0.5 ng/mL).

#### 1.7 Imaging acquisition

For nuclear counterstaining, fixed colonies were incubated with DAPI (Sigma Aldrich; 1:1000 in PBS) for 90 min at room temperature, followed by three washes in PBS. Patterned chips were then mounted on glass slides using Fluoromount-G (Thermo Fisher) to preserve fluorescence. Imaging was performed using Zeiss LSM-800 confocal laser scanning microscope equipped with a 20× objective.

### 2. Materials and Methods: Theory and modeling

#### 2.1 Defining Order parameter (𝒪_p_)

We consider a colony of fixed size *N* cells. For colony *m*, let *x*_*m*j_ ∈ {0,1} denote the binary fate of cell *j* (e.g., SOX2-high = 1, otherwise 0). The colony composition, *i* (*i* ∈ {0,1, …, *N*}) is defined as the number cells with the states *x*_*m*j_ = 1 in the colony:

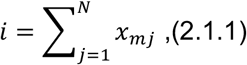

##### Macro-states and probabilities

Across many colonies with the same number of cells *N*, we define *P*_*i*_ as the probability that a randomly chosen colony has the composition *i. P*_*i*_ defines the macro-state distribution and it satisfies 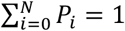 and 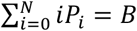.where *B* is the expected number of cells with state 1 (SOX2-high cells) per colony.

##### Micro-states and entropy

Each colony composition *i* corresponds to many distinct microstates (specific cell-level configurations). The number of such microstates is 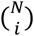. Since we consider cells with identical state in the colonies identical, every microstate with the same composition *i* is equally likely, with the microstate probability of:

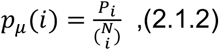

The microstate entropy is given by(*52*):

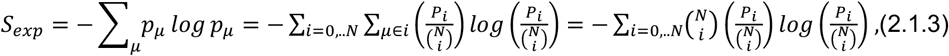

Therefore:

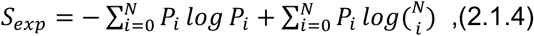

This quantity measures the observed disorder across all possible cell-level states, given the empirical distribution of compositions (*P*_*i*_).

##### Maximum entropy reference

To interpret the level of order, we compare to the maximum possible entropy under the same constraints(*53*). We will maximize the microstate Shannon entropy, while accounting for the degeneracy of macrostates 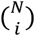.

To do so, the Lagrangian function ℒ is:

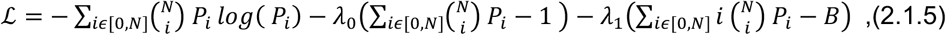

Next, we find the extremum value with respect to probability distributions (P_i_) by setting:

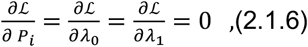

Stationarity gives:

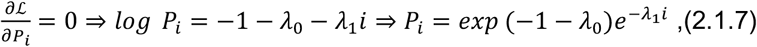

Replacing this into the 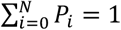, yields:

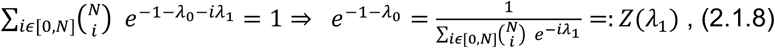

Normalization yields a “partition function” and therefore:

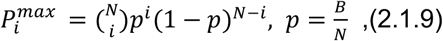

This corresponds to the case where cells act independently, with probability *p* of being state 1, but with the same expected mean *B*. The corresponding maximum entropy is:

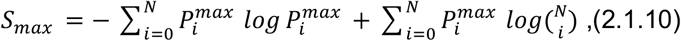

##### Order parameter

We define the order parameter as the normalized entropy gap:

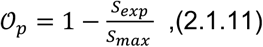

Where 𝒪_*p*_ → 0 corresponds to maximally disordered systems, 𝒪_*p*_ → 1 corresponds to highly ordered systems, and 0 << 𝒪_*p*_ << 1 corresponds to intermediate levels of order (see Supplementary Methods: Analysis for details).

#### 2.2 Single-cell Bistable switch model

We modeled the SOX2-high/CDX2-low vs SOX2-low/CDX2-high decision as a toggle switch with mutual repression and first-order decay(*29*). After nondimensionalization, the core (single-cell) system of ordinary differential equations (ODEs) is (**Fig. S9 a**):

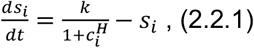

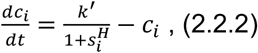

Where *s*_*i*_ and *c*_*i*_ denote SOX2-high/CDX2-low and SOX2-low/CDX2-high states for cell *i. H* is the Hill coefficient (default *H* = 2), and *k, k′* set maximal (dimensionless) production rates. We assumed the same decay rate for both species for simplicity. Characterizing the behavior of the system of ODEs based on (*k, k′*) yields a bistable region and a monostable region (**Fig. S9 b**). For all core analyses, we used the symmetric case (*k*=*k′*) within the bistable domain (default *k* = 3). Simulations of the model to steady state used the Runge-Kutta method of order 4. To capture intrinsic variability, we extended the deterministic toggle with small, independent Gaussian fluctuations on production terms (*σ dW*(*t*)), where *dW*(*t*) is independent Wiener increments. We used *σ* = 0.7 and Euler-Maruyama to solve the resulting stochastic differential equations. To probe single-cell dynamics, simulations were initialized with *c*_*i*_ = 0 (*s*_*i*_ only state). To destabilize the system and reveal both attractor basins, we applied a transient square-pulse increase in the production of *c*_*i*_.

#### 2.3 Investigating possible signaling configuration topologies

We extended the single-cell toggle to a two-cell system with intercellular non-paradoxical signaling (as opposed to paradoxical signaling(*54, 55*)). Given the size of our system and movement of the cells, we assumed that signaling is well-mixed. Each cell *i* ∈ {1,2} carries two intracellular states *s*_*i*_ and *c*_*i*_ and secretes two extracellular ligands, *w* (secreted by *s*_*i*_) and *z* (secreted by *c*_*i*_) that mediate communication across cells.

The intracellular dynamics are:

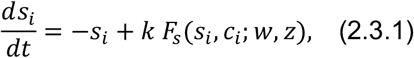

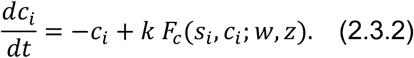

Regulation is implemented using a pooled Hill function(*56, 57*):

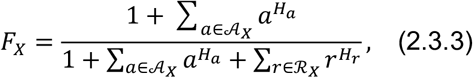

Where *X* ∈ {*s, c*}. Here, 𝒜_*X*_ and ℛ_*X*_ denote the sets that contain activators and repressors for target *X*, with Hill exponents *H*_*a*_, *H*_*r*_ set to 4 for intracellular repressors and 1 for ligand-mediated interactions. The core toggle is always present (*c*_*i*_ ∈ ℛ_*s*_, *s*_*i*_ ∈ ℛ_*c*_). Additional edges are optional and are controlled by four topology flags. Each flag determines whether a secreted ligand acts as an activator, a repressor, or is absent:

*Flag*_*s,w*_: Self-regulation of *s* via *w*:

if flag is “+”: *w* is added to the activator set 𝒜_*s*_

if flag is “-”: *w* is added to the repressor set *R*_*s*_.

if flag is “∅”: *w* is not added.

*Flag*_*c,z*_: Self-regulation of *c* via *z*:

if flag is “+”: *z* is added to the activator set 𝒜_*c*_

if flag is “-”: *z* is added to the repressor set *R*_*c*_.

if flag is “∅”: *z* is not added.

*Flag*_*s,z*_: Cross-regulation of *s* via *z*:

if flag is “+”: *z* is added to the activator set 𝒜_*s*_

if flag is “-”: *z* is added to the repressor set *R*_*s*_.

if flag is “∅”: *z* is not added.

*Flag*_*c,w*_: Cross-regulation of *c* via *w*:

if flag is “+”: *w* is added to the activator set 𝒜_*c*_

if flag is “-”: *w* is added to the repressor set *R*_*c*_.

if flag is “∅”: *w* is not added.

Each flag can take one of three values, absent (∅), activating (+), or inhibitory (-), yielding 3^4^=81 possible signaling topologies. The extracellular ligands are shared across the two cells, with production and decay given by:

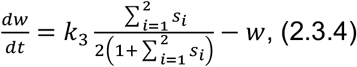

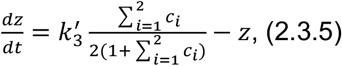

For consistency, we scanned signaling strengths 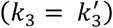 in the range of [1,2]. This range was chosen to ensure that the ligand level could bias the gene expression state of the bistable switch, while maintaining bistability (**Fig. S9 c** and **d**).

#### 2.4 Multicellular model and signal dilution

We extend the single-cell toggle to an *n*-cell colony communicating via soluble ligands. The key effect of colony size is how secreted signals diffuse into the available space (**Fig. S11 a**), which we assume is proportional to the colony’s extra-nuclear volume(*30, 58*). We model how this available ‘reservoir’ changes with the number of cells on the adhesive island and use it to adjust the “effective ligand concentrations” accordingly. We assume tissues are effectively 2D monolayers.

##### A) Mechanical packing model for the extra-nuclear reservoir

To build a mechanical model, we first look at cells that are unconfined. In this state, the available area for a cell to spread freely is more than its need and each relaxed (unconfined) cell has cross-sectional area of:

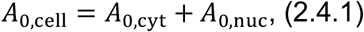

Where *A*_0,cyt_ and *A*_0,nuc_ are extra-nuclear and nuclear contributions. When *N* cells are placed on a circular pattern with radius *R*, two regimes follow area constraint of the pattern:

i. *NA*_0,cell_ ≤ *πR*^2^: The cells are not mechanically under stress by limited space, therefore:

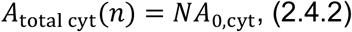
ii. *NA*_0,cell_ > *πR*^2^: The cells are confined (**Fig. S11 b**), and under mechanical constraint. In this case, nuclear and cytoplasmic areal strains share the required compression elastically. We define strains as:

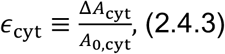

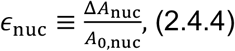

Where Δ*A* > 0 denotes area compression. Therefore, the per-cell post-compression areas for nuclear and cytoplasmic parts are:

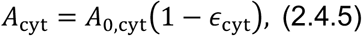

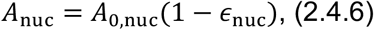

We assume linear areal elasticity with effective moduli *K*_cyt_, *K*_nuc_. We can determine the strains by energy minimization under the packing constraint. The areal packing constraint that denotes the total area covered on an adhesive island should be equal to the total available area:

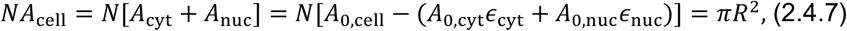

The elastic energy is(*59, 60*):

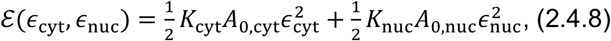

Minimizing elastic energy (based on strains) yields the stationarity conditions:

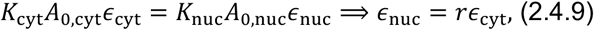

Where 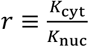.

Combining with the packing constraint gives:

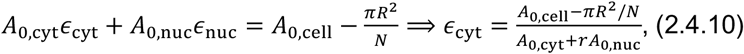

Thus, the per-cell cytoplasmic area under confinement is:

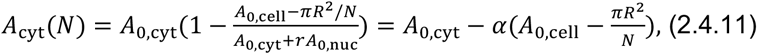

Where 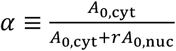.

Putting both regimes together, the total cytoplasmic area is:

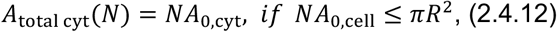

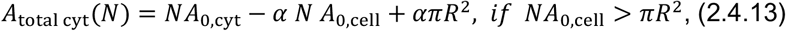

##### B) 𝒦 (dimensionless dilution factor, compactness)

We define 𝒦 as:

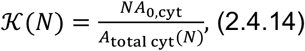

Therefore, (*N*) = 1 in the unconfined regime, and 𝒦(*N*) > 1 once confinement reduces per-cell cytoplasmic area. The value of *R* = 28 (raw pixels, equal to 35 um) is measured from imaging. *A*_0,cyt_ = 173 ± 51 and *A*_0,nuc_ = 64 ± 20 are determined from imaging of patterns with only one cell. The value of *r* varies in the literature based on technique and cell differentiation stage, but ranges from 0.1 to 0.9(*61–64*). For our study we assumed *r* = 0.5 as the middle value. Evaluating the model with different values of parameters in the range of our measurements of *R, A*_0,cyt_ and *A*_0,nuc_ (values drawn from a distribution) showed that *A*_total cyt_(*N*) start to increase linearly with number of cells until the systems reaches the mechanically constrained phase. After that, *A*_total cyt_(*N*) starts to either increase with a smaller slope or saturates (**Fig. S11 c**). Importantly, experimental measurements of 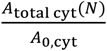 support the model predictions (**Fig. S11 d**).

###### Multicellular signaling ODEs with signal dilution

For a multicellular system with *N*cells (*i* = 1, . . *N*) with *s*_*i*_ and *c*_*i*_ as the toggle states and *w* and *z* as the two shared ligands (with background *z* ligand of *b*_0_), the concentration dynamics of ligands are:

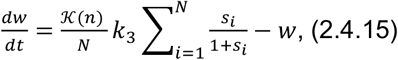

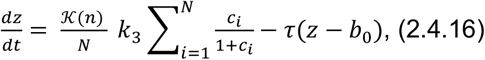

The free parameter of the final model (*k*_3_) has been fitted to the experimental order parameter readouts (from **Fig. S7 b**) for adhesive islands with radius of 35*µm* (**Fig. S12 a** and **b**).

#### 2.5 Modeling the effect of Guide-cell

To investigate the ability of rare “guide” cells to influence fate choice in confined colonies, we extended the multicellular framework to include sender-receiver interactions. Each cell carries the same endogenous bistability. However, a small number of cells in each colony (*n*_*guide*_ = 1) were designated as sender cells (guide cells), capable of producing exogenous BMP4 upon induction with doxycycline (DOX). Sender secretion was modeled as an additional ligand input into the extracellular pool of ligands.

The coupled system of ODEs was integrated for colonies of size *N*_*cell*_ = 2 − 20, with a single guide-cell (*n*_*guide*_ = 1) and the remainder were normal cells (receivers). For each colony, the cytoplasmic dilution model was used to scale the effective extracellular ligand concentrations. Guide cells secretion was modeled as an additive source term in the extracellular ligand pool. Thus, in addition to endogenous secretion of *z*, the guide cell contributes a constant secretion rate of γ:

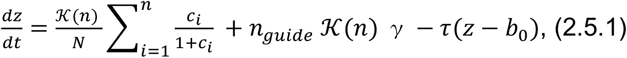

Where (*n*) is given by equation 2.4.14.

### 3. Supplementary Text: Analysis

#### 3.1 Cell segmentation and gene readout binarization

Colonies were imaged on a Zeiss LSM-800 confocal microscope with a 20X objective. Channels were acquired for DAPI (nuclei), and SOX2. Images were processed in CellProfiler (v5)(*65*) using a standardized pipeline. Briefly, after a mild Gaussian smoothing, nuclei compartments were segmented from the DAPI channel. Individual cells were identified using adaptive thresholding and de-clumping, and then merged into colony objects based on proximity. Per-cell mean and integrated intensities were measured for each channel, and colony membership was recorded. The parental colony identity was used for colony level analysis later (**Fig. S4a**).

For downstream analysis, SOX2 intensities were normalized per image to control for exposure differences, and then binarized into positive/negative states. Thresholds were determined from the distribution of intensities in each experiment, using k-means clustering (k =2 for two clusters), (**Fig. S4b**). This produced consistent binary classifications of cell states. Colony-level features such as size and fate composition were then computed by grouping cells according to their parent colony ID. This workflow ensured reproducible extraction of single-cell states and colony structure, enabling quantitative analysis of fate distributions and comparison with theoretical models. Further downstream calculations were performed in customized Python-based workflows.

#### 3.2 Evaluation of the Order Parameter (𝒪_p_) using synthetic data

To systematically assess how the order parameter (𝒪_p_) reflects multicellular organization, we constructed synthetic ensembles of colonies manually in ordered form, and consequently injected disorder into the synthetic ensembles and investigated the effect of added randomness on 𝒪_p_ (**Fig. S6a**). Each synthetic colony consisted of *N*_*cell*_ cells, each in a binary state (0 or 1). The ensemble (which consists of *N*_*colony*_ colonies) was initialized in the most ordered configuration to maximize colonies which are entirely composed of state {0} or state {1} cells. We made sure that the overall population (*N*_*cell total*=_*N*_*colony*_ × *N*_*cell*_) fractions matched the prescribed probability of cell state distribution, *P*_0_ + *P*_1_ = 1 (**Fig. S6a**).

To progressively introduce disorder, we performed cross-colony shuffling. At each shuffle step, two different colonies were chosen at random, and one cell from each was swapped provided that their states differed. This procedure preserves the global state fractions while redistributing cells across colonies. The number of accepted swaps therefore acts as a tunable disorder parameter. For each colony *N*_*cell*_ and number of colonies *N*_*colony*_ we computed 𝒪_p_ at successive shuffling levels using bootstrap resampling (with sampling parameters consistent with the calculations for experiments and models) to estimate variability. Across all colony sizes, 𝒪_p_ decayed monotonically with increasing shuffle count, consistent with a loss of collective order (**Fig. S6b**). Importantly, the relevant control variable is not the raw number of swaps, but their density relative to the total number of cells (*N*_*cell total*_). Normalizing the horizontal axis by the total number of cells, this yields:

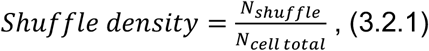

This renormalization caused the 𝒪_p_ curve for different colony sizes to collapse onto a single master curve. This collapse demonstrates that 𝒪_p_ consistently captures the degree of order across systems of different sizes (**Fig. S6c**), and different values of cell state distributions (*P*_0_, *P*_1_) (**Fig. S6d**).

#### 3.3 Calculating Order Parameter (𝒪_p_) for samples

To quantify the degree of coordination in fate decisions across cells within each colony, we computed an entropy-based order parameter (𝒪_p_) capturing deviation from an independent, uncoordinated model. This was applied to both experimental and simulated data where each cell was binarized as either SOX2+/CDX2- or SOX2-/CDX2+ (or equivalently, state 1 or 0). We calculated 𝒪_p_ as defined before:

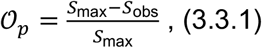

To assess the robustness and variance of 𝒪_p_ across colony sizes, we performed bootstrap sampling of experimental colonies. For each of *N* = 100 bootstrap runs, 60% of colonies were sampled (with replacement), and the 𝒪_p_ was computed independently per condition and colony size. Within each bootstrap replicate, we excluded colonies with < 5 cells. For colonies with one cell, coordination is undefined and 𝒪_p_ was set to zero. To summarize these distributions, we computed the mean and standard deviation of *𝒪*_p_ per colony size across bootstrap replicates. For each size we have 3 experimental replicates (separate chips, see **Fig. S7a** and **b** for individual replicate breakdowns). The final reported 𝒪_p_ values have been based on pooling all 3 replicates together (**Fig. S7c**). To identify the trend, we applied a sliding smoothing window to collect the mean, with window size 3 centered at each colony size *n*. For each window, we computed:

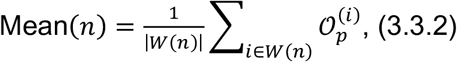

Where *W*(*n*) is the set of 𝒪_p_ values in the window centered at size *n*.

#### 3.4 scRNA-seq analysis

##### Preprocessing and Quality Control

Single-cell RNA sequencing data were analyzed using the Scanpy Python package (v1.10.2)(*66*). Initial quality control steps were applied to filter out low-quality cells and outliers. Cells were retained if their total UMI count was between 20,000 and 77,000. Summary metrics, including the number of detected genes per cell, total counts per cell, and the percentage of mitochondrial transcripts, were visualized via violin and histogram plots to inform thresholding (**Fig. S3a-e**). Cells outside the defined UMI range were excluded from downstream analysis (**Fig. S3f**).

##### Normalization and Highly Variable Gene Selection

Raw count matrices were stored in an Anndata object for downstream reproducibility. Cell-wise normalization was performed by scaling each cell’s total counts to a common median library size, followed by logarithmic transformation. Highly variable genes (HVGs) were identified using Scanpy’s default method, selecting the top 2,000 HVGs to capture the most informative transcriptional features for dimensionality reduction and clustering (**Fig. S3g** and **h**).

##### Dimensionality Reduction and Clustering

Dimensionality reduction was performed using principal component analysis (PCA) on the log-normalized expression of HVGs. A neighborhood graph (*number*_*of*_*neighbors* = 50) was computed based on the first 50 principal components. Clustering was then performed using the Leiden algorithm(*67*) with a resolution parameter of 0.6 followed by breaking up a wide-spread cluster into 3 clusters (resolution parameter of 0.3) that yielded clusters C3, C6 and C8 (**Fig. S3i** and **j**).

##### Cluster Annotation and Marker Gene Analysis

Clusters were annotated using known lineage-specific marker genes(*68, 69*). Gene sets included pluripotency markers (SOX2, POU5F1, NANOG, MYC, and DPPA4), ectodermal markers (PAX6, SOX1, FOXJ3, OTX2, and IRX3), extraembryonic markers (CDX2, GATA3, TFAP2A, HAND1, and EPCAM), mesodermal markers (TBXT, EOMES, MESP1, SNAI1, and PDGFRA), and endodermal markers (SOX17, HNF1B, FOXA2, GATA6, and CXCR4). Expression of these markers was visualized using dot plots and PCA overlays (**Fig. S3k** and **l**). Differential expression analysis was performed using the Wilcoxon rank-sum test and top-ranking genes per cluster were visualized using dot plots and summary figures to assist annotation.

##### Cluster Evaluation Using Marker Gene Scores

To evaluate the biological coherence of clusters, a custom scoring strategy was employed. For each annotated marker gene group, z-scored expression values were computed per cell and averaged within each cluster. In parallel, the fraction of cells within each cluster expressing genes above a defined threshold was also computed. These metrics were combined to generate a dot plot (**Fig. S3m**), where color represented the mean scaled expression and dot size indicated the proportion of cells expressing the gene set in the main text. This approach provided a high-level and quantitative evaluation of cluster identity. Clusters were renamed accordingly based on combined marker expression and time point.

##### Timepoint Composition Analysis

To evaluate how cluster composition changed over time, each cell was assigned a timepoint corresponding to the sample ID (0, 12, 24, or 36 hours after BMP4 stimulation). The fraction of cells from each timepoint was computed for every Leiden cluster. These values were visualized using a stacked bar plot to assess temporal shifts in cluster occupancy, revealing dynamic population transitions in response to BMP4 exposure (**Fig. S3o** and **p**).

##### Expression Visualization on SOX2-CDX2 axis

To reduce technical noise and resolve joint gene expression relationships, MAGIC-based imputation (v3.0.0) was applied using a minimal imputation setting (*knn* = 2). This was implemented via a custom wrapper function storing the resulting imputed values in an adata layers. All genes were used for imputation. Next, the imputed expression of SOX2 and CDX2 was visualized using scatter plots, with cells colored by annotated group identity. These visualizations helped resolve distinct subpopulations based on the joint distribution of fate-associated markers.

##### Kernel Density on Marker Expression

Visualization (SOX2-CDX2 Axis) To examine dynamic changes in marker expression distribution over time, imputed SOX2 and CDX2 expression values were extracted per cell and plotted using 2D scatterplots overlaid with kernel density estimate (KDE) contours. Separate plots were generated for each timepoint (0, 12, 24, and 36 hours post-BMP4 stimulation), with contours indicating local cell density in expression space. These plots revealed a progressive shift in SOX2-CDX2 distribution and bifurcation, consistent with cell state transitions (**Fig. S3q**). Additionally, we have quantified the endogenous BMP4 expression via clusters. Analysis shows a clear separation between clusters 4, 5, and 6 (marked as SOX2high/CDX2 low at 36 h) vs. 7 and 8 (marked as SOX2low/CDX2 high at 36 h), (**Fig. S3r**).

#### 3.5 Cell-Cell Communication and Ligand-Receptor-TF path classification

##### Ligand receptor interaction database

To investigate how transcriptionally distinct cell populations influence one another through signaling, we developed a custom multi-layered framework that integrates ligand-receptor inference, cluster-specific ligand expression, and receptor-TF correlation. The pipeline combined CellChat’s ligand-receptor interaction prediction with a custom receptor-TF scoring system.

We began by applying the CellChat(*27*) R package (v1) to the processed single-cell transcriptomic data using the curated human interaction database. Cells were grouped using the biologically annotated states such as “SOX high CDX2 low” and “SOX low CDX2 high”. CellChat’s built-in functions were used to identify overexpressed ligands and receptors within each group, compute potential ligand-receptor interactions, and project expression data onto a protein-protein interaction network to account for known physical and functional associations. To ensure robustness, we filtered out any inferred communication events in which either the source or target cluster had fewer than 10 contributing cells. The resulting interaction predictions were extracted from the CellChat object for use in next steps.

##### Mapping UniProt Interactions to Gene Symbols

To align the protein-level interaction data from CellChat with gene-level expression matrices, we parsed metadata from the CellPhoneDB (*70*) database, including protein, complex, and gene tables. For each interaction, the two components were parsed to determine their molecular identity (ligand or receptor) and gene content. Both proteins and complexes were resolved into lists of UniProt IDs, which were then mapped to gene names using a curated lookup table. This process yielded a complete gene-level interaction map annotated with ligand and receptor roles.

##### Ligand usage matrix, *M*_1_

We then organized these gene-level interactions into a nested dictionary indexed by source and target clusters. A custom query function was used to retrieve, all participating ligands, receptors, or ligand-receptor pairs. This enabled the construction of a ligand usage matrix *M*_1_ ∈ ℝ^*S*×*L*^, where *S* denotes the set of source clusters and *L* the set of ligands. For each source cluster *S*, differential expression analysis was performed using the Wilcoxon rank-sum test implemented in scanpy. Ligand genes were retained only if their log fold-change was positive, applying the threshold:

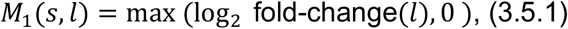

Downregulated ligands (log_2_ fold-change ≤ 0) were explicitly set to zero, effectively excluding them from further analysis (**Fig. S8a**). The resulting matrix *M*_1_ was then normalized across all entries to the range [0,1] to yield a relative expression enrichment score for each ligand per cluster (**Fig. S8b**).

##### Ligand-receptor adjacency matrix, *M*_2_

To model molecular interaction logic, we constructed a binary ligand-receptor adjacency matrix *M*_2_ ∈ {0,1}^*L*×*R*^, with rows representing ligands and columns representing receptors. An entry *M*_2_(*l, r*) = 1 indicated that ligand *l* and receptor *r* form a known functional pair based on the database, while a zero indicated no direct interaction (**Fig. S8c**).

##### Receptor–TF correlation matrix *M*_3_

To estimate the influence of receptors on target transcription factors, we defined a third matrix *M*_3_ ∈ ℝ^*R*×*T*^, where *R* is the set of receptors and *T* the set of transcriptional states corresponding to target clusters. To build this matrix, we first imputed gene expression using MAGIC (v3.0.0) with a conservative setting (knn = 2) to avoid over-smoothing. For each receptor-TF pair (*r, t*), we calculated the Pearson correlation coefficient:

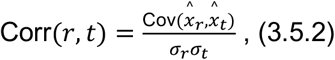

where 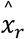 and 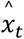 are the imputed expression vectors of receptor and TF. We additionally calculated conditional Mutual information(*71*) in order to capture nonlinear dependencies:

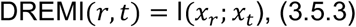

The Receptor-TF correlation matrix *M*_3_ is defined as:

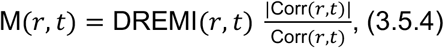

In visualization and network construction steps, we applied an additional filter, retaining only receptor-TF pairs with |Corr(*r, t*)| > 0.5 (**Fig. S8d**).

##### Path Score

With all three matrices defined, we computed a signaling score for each possible four-node pathway connecting a source cluster *s* to a target cluster *t* via ligand *l* and receptor *r*:

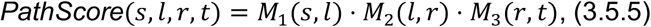

Only paths with non-zero scores (PathScore ≠ 0) were retained for ranking and visualization. This product formulation enabled the integration of upstream expression, molecular feasibility, and downstream correlation into a single signaling inference model.

##### Visualization

We visualized top-scoring ligand-receptor-TF triplets for each cluster pair using ranked scatter plots. Ligands were colored by their associated signaling pathways (e.g., BMP, WNT) as annotated in the CellPhoneDB database. Pathways were further grouped, and their corresponding ligands and receptors were assigned consistent colors to enhance visual clarity. To reveal the full communication topology, we constructed a directed four-layer network using NetworkX. Nodes represented source clusters, ligands, receptors, and target clusters, respectively. Edges between nodes were derived from the *M*_1,2,3_. Specifically, source-to-ligand edges reflected differential expression (gray scale from *M*_1_), ligand-to-receptor edges reflected binary interactions (colored by ligand pathway from *M*_2_), and receptor-to-target edges reflected receptor-TF correlations (blue-red scale from *M*_3_).

To summarize the flow of communication across the entire system, we grouped all valid paths by source and target cluster and counted the number of paths with positive or negative scores. These counts were visualized using Sankey diagrams, with nodes on the left representing source clusters and nodes on the right representing target clusters (**Fig. S8e** and **f**). Link widths were proportional to the number of high-confidence signaling paths, and link colors indicated the net directionality of influence: blue for positive, red for negative. Only paths with non-zero scores were included. Together, this framework provided a scalable and interpretable approach for linking extracellular signaling events to lineage-informative transcriptional programs, revealing possible paths of information transfer between cells.

#### 3.6 Inter-colony Spatial Correlation Analysis via Moran’s I

To assess spatial coordination of cell fate across colonies, we quantified inter-colony spatial autocorrelation in SOX2 expression using Moran’s I statistic(*72*). Each colony (i) was segmented using CellProfiler, and its spatial position was defined by the centroid of its bounding region. The fraction of SOX2-positive cells per colony was computed using a mean intensity threshold and binarization step defined upstream in the segmentation pipeline. We constructed spatial graphs between colonies based on radius (bandwidth) distances. Spatial weight matrices *W* were constructed for multiple thresholds *d*, with colony pairs connected if their centroids were within Euclidean distance *d*. In all cases, the spatial weights were row-normalized to ensure comparability across thresholds. The spatial autocorrelation of SOX2-positive cell fractions ***x*** across colonies was then quantified by Moran’s I, defined as:

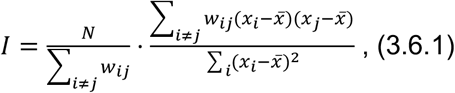

Where *N* is the number of colonies, *w*_*ij*_ is the spatial weight between colony *i* and *j*, and 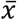 is the global mean of the SOX2-positive fraction cells across all colonies in the neighborhood domain. To ensure the robustness, we picked the neighborhood domain in a range of values to include statistically 5 or more neighbors (**Fig. S5a** and **b**). Significance was assessed using permutation tests. For each spatial weight matrix, we performed 999 random permutations of the observed colony SOX2 fractions and computed Moran’s I for each shuffled configuration. The empirical p-value was calculated as the proportion of permutations yielding a Moran’s I greater than or equal to the observed value (**Fig. S5c**).

#### 3.7 Clonal analysis

##### Calculation

Within each colony we regressed normalized SOX2 intensity on normalized *CellTracker* intensity using linear regression which gives slope, intercept, Pearson *r*, and a two-sided t-test p-value for non-zero slope. We computed 95% CIs for the mean prediction along the fitted line using the standard OLS formula with Student-t quantiles (*n* − 2).

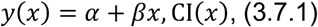

##### Visualization

For each colony we plotted the scatter (SOX2 vs *CellTracker*), the regression line, and its 95% CI ribbon. Lines were colored red for significant positive *r*, blue for significant negative *r*, and light grey otherwise. A summary “volcano” displayed colony-wise *r* versus −log_10_ (*p*) with a horizontal line at *p* = 0.01 and a vertical line at *r* = 0; points were colored as above. Percentages reflect the fraction of colonies in each category.

#### 3.8 Categorization and scoring the results of the search on signaling configurations

For each topology *T*, we simulated the two-cell system from 100 random initial conditions and classified the steady-state outcome according to the relative dominance of *s*_*i*_ and *c*_*i*_ in each cell. We called a two-cell state:

i. SS: if s_1_ >> c_1_ and s_2_ >> c_2_,
ii. CC: if s_1_ << c_1_ and s_2_ << c_2_,
iii. SC: if one cell was s s-dominant and the other c c-dominant.

This yielded empirical probabilities *P*_*SS*_(*T*), *P*_*CC*_(*T*) and *P*_*SC*_(*T*) for each two-cell state of the signaling topology *T* (**Fig. S10a-c**). From these, we computed three normalized metrics (**Fig. S10d**):

1. Minimality (favoring parsimonious topologies):

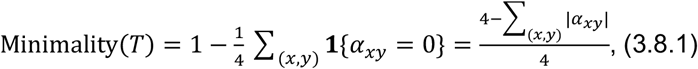 Where *α*_*xy*_ are possible inter-cellular interactions. Note that if all possible interactions are present Σ_(*x,y*)_ |*α*_*xy*_|= 4, and if non is present, Σ_(*x,y*)_ |*α*_*xy*_| = 0.
2. Cohesion (propensity for consensus outcomes):

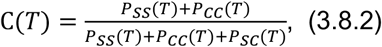
3. Diversity (balance between the two consensus fates):

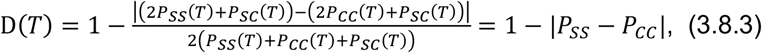

We used the fact that *P*_*SS*_ + *P*_*CC*_ + *P*_*SC*_ = 1, to further reduces the expression of *D*. A composite score was defined as the weighted geometric mean of the three metrics (**Fig. S10d**) in a way that minimality score does not dominate. This scoring scheme allowed us to rank all possible signaling configurations by their parsimony, stability, and fate-balance properties (**Fig. S10e**) and selecting the best possible set of interactions (**Fig. S10f**).

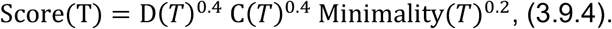

#### 3.9 investigating the effect of cell division on *𝒪*_p_

To examine how proliferation influences collective order, we implemented a multicellular model in Morpheus software(*73*) that explicitly tracks both cell states and cell division

The equations of state and environment are similar to the original model. Colonies were simulated on a two-dimensional lattice using the Cellular Potts framework, with each cell represented as a contiguous domain, subject to volume and surface constraints. Cells carried internal state variable (*s*_*i*_ and *c*_*i*_ for the *i*^*th*^ *cell*). The colony dynamics were evolved for simulated time of 42h, with one Monte Carlo Step (MCS) corresponding to one minute of biological time. Colonies were initialized with cells placed randomly within a circular region approximating the micropattern radius, and allowed a short equilibration before state updates commenced.

Cell division was incorporated using Morpheus CellDivision plugin. Each cell carried a cycle clock, which advanced with each MCS. At birth, cells were assigned a cycle length drawn from an normal distribution with mean 900 minutes, corresponding to a 15-hour average doubling time measured for human pluripotent stem cells(*74, 75*). Once the cycle clock exceeded the assigned cycle length, the cell divided into two daughters. The parental lattice domain was split into two daughter compartments, each with half the original target volume and a small positional displacement to avoid overlap. Daughters inherited the cellular state of the parent. At the conclusion of each simulation, the cell state composition of every colony was recorded, and the order parameter *𝒪*_p_ was computed exactly as defined previously.

To dissect the effect of cell division vs. the effect of cell-cell communication, this procedure was applied under four regimes:

a. No cell-cell communication and no cell division.
b. Cell-cell communication without division.
c. No cell-cell communication and cell division.
d. Both cell-cell communication and cell division.

In the absence of both processes, colonies exhibited low order, with *𝒪*_p_ values near those expected for independent cells. Enabling communication increased order and followed the overall trend. Enabling division increased order by amplifying stochastic imbalances in small cell numbers. When both communication and division were present, the two effects reinforced one another, yielding the highest *𝒪*_p_ across colony sizes (**Fig. S13**). In summary, the presence of cell division only caused a mild shift in the order parameter of colonies with a small number of cells.

#### 3.10 Analyzing information propagation via induction factor

##### Definition of induction factor

To quantify how the information of perturbation in the state of one cell (the “inductor”) propagates into the shared environment, we measure a field-level induction factor, χ, that captures the effect of the inductor’s change of state on the shared environment. In each simulation, we apply a finite pulse *Pulse*(*t*) to the inductor’s production of the species c between *T*_0_ and *T*_0_ + *T*_dur_ as (**Fig. S14a** and **b**):

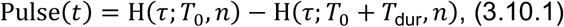

where H is a smoothed step function:

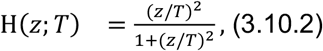

We evaluated the time-averaged states in two windows that bracket this pulse. Field displacement is computed as the Euclidean distance between the signaling states (*E* = (*w, z*)) of the system before and after the pulse as:

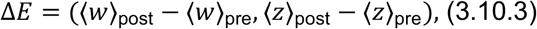

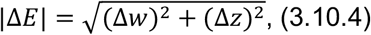

To normalize input strength, we use the inductor’s own move in gene space over the same windows:

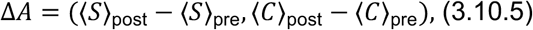

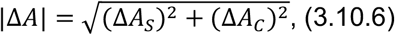

And define the induction factor (similar to susceptibility(*76*)) as (**Fig. S14b**):

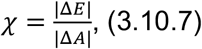

##### Fitting a bi-sigmoid curve to find trends

To summarize how field induction scales with colony size *N*, we fit the replicate-level χ(*N*) with a bi-sigmoid(*77*) curve by non-linear least squares:

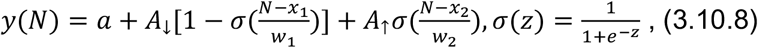

Here *a* is the baseline, *x*_1_ and *x*_2_ are transition locations, *w*_1_ and *w*_2_ control steepness, and *A*_↓_ and *A*_↑_ set step amplitudes of descent and ascent. We initialize parameters from empirical quantiles and constrain widths and locations to the observed *N* range. We perform this analysis for all three different colony radii (**Fig. S14c**).

#### 3.11 Kullback–Leibler (KL) divergence analysis of gene expression distributions (colonies with vs without a guide cell)

##### Cell and Colony Filtering

To quantify how guide cells alter SOX2 expression, we computed the KL divergence(*38*) between GFP (SOX2) distributions in normal (BFP^+^) cells across colony sizes, comparing colonies with vs. without a single guide cell (BFP^−^/GFP^−^). We selected colonies containing exactly 0 or 1 guide cells (BFP^−^) and analyzed only the BFP^+^ population (receiver cells). Colony size was computed and binned into 5 groups (1-20 cells).

##### Distribution Estimation

For each colony size bin and condition, we constructed histograms of normalized SOX2 (GFP) intensity for BFP^+^ cells, grouped by presence or absence of a guide cell. Histograms used 6 GFP bins over [0, 1], and were conditionally normalized within each size bin to yield probability distributions.

##### KL Divergence

For each size bin, we computed KL divergence between GFP distributions in colonies with vs. without a guide cell:

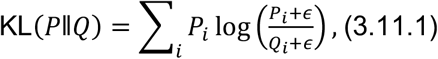

where *P* is the distribution of colonies with one guide cell, and *Q* is the distribution of colonies with no guide cell. To ensure numerical stability, *ϵ* = 10^−9^ is added. This metric quantifies shifts in SOX2 expression attributable to guide cell presence. KL values were visualized as heatmaps and saved alongside the raw histograms for each condition. Results highlight how a single guide cell can modulate SOX2 expression in neighboring cells depending on colony size.

##### Bootstrap uncertainty

To obtain uncertainty and robust summaries per size bin, we performed bootstrap resampling (n=20). For each bin, we first down-sampled both groups to equal size (without replacement down to 70% of original sample sizes) and then drew bootstrap resamples from each group to recompute KL divergence (**Fig. S15**).

#### 3.12 Connecting Guide cell experiment to model

##### A simple model for DOX inducible BMP4-RFP secretion

We model the number of active construct copies per cell as a Poisson random variable(*78*).

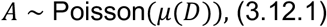

With the mean of:

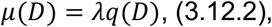

Here *λ* is a constant and *q*(*D*) is the probability that each construct can be activated, which can be modeled as a Hill function(*79*):

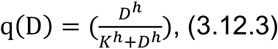

Where D is the Dox concentration in the media, and K is a constant. The probability that a cell be activated (ON) - which means the cell has at least one active construct- is given by:

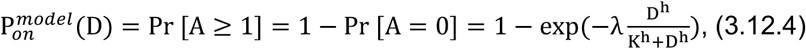

##### Connecting to the flow cytometry measurement of RFP

To measure the cell in the ON state experimentally, we define a fixed RED-channel threshold T. the value of T is driven from WT cells in a way that WT cells (cells with no transduction) be filtered out as the background. We set T at a chosen high percentile of the WT distribution (95%). For a given dose D, we define the Fraction ON as the ratio of cells that has their RFP bigger than the threshold, to all cells:

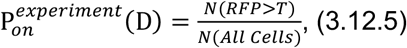

Comparing equation (3.12.4) and experimental measurements from equation (3.12.4), yields the best fit of values 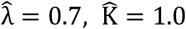, and 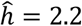. The curve was fitted to the ON fraction readouts (**Fig. S17a**). To test and validate our model, we plotted the whole distribution of predicted RFP intensities, for a range of DOX concentrations. The results showed that our simple Poisson+Hill model can reproduce the distributions (**Fig. S17b**). We have noticed a potential reduction in RFP in DOX concentrations greater than 2 ug/ml, that could be potentially due to toxicity.

##### Anchoring the signal secretion of the guide cell model (*γ*) to the mean values of RFP

Experimentally FACS sorted cells were modeled as cells with their RFP signal being higher than threshold value, T. The RFP threshold value was set to the 95th percentile of RFP value of the WT cells. For each DOX concentration, D, we computed the mean of RFP readouts of the selected cells, *μ*_*RFP*_ (*D*).

Next, to calibrate the guide-cell model, we used a single experimental condition DOX = 1 µg ml^−1^ and only used the bin of 16-20-cells. For this anchor (DOX = 1 µg ml^−1^, bin of 16-20), we extracted the experimental guidance values (KL divergences) and compared them to the guide-cell simulations evaluated across the grid of secretion factors (γ) at the same bin (16-20-cells). This yielded γ =γ* as the best fitted secretion factor to the experimental guidance at this condition. Here we only looked at the mean-field behavior of the model for simplicity. Assuming that BMP4 secretion of cells is proportional to RFP readouts (note the inducible expression of BMP4-2A-RFP), we can define this relationship between the best fitted (BMP4) secretion factor, and the measured RFP (**Fig. S17d**):

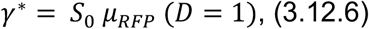

Where S_0_ is a positive scalar. This establishes a correspondence between flow RFP readouts and the model’s secretion factors γ for other experimental conditions (Dox=D), (**Fig. S17d**):

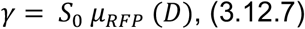

and allows us to directly compare the guidance of the guide-cell model and experiments.

### Supplementary Figures

**Fig. S1.**
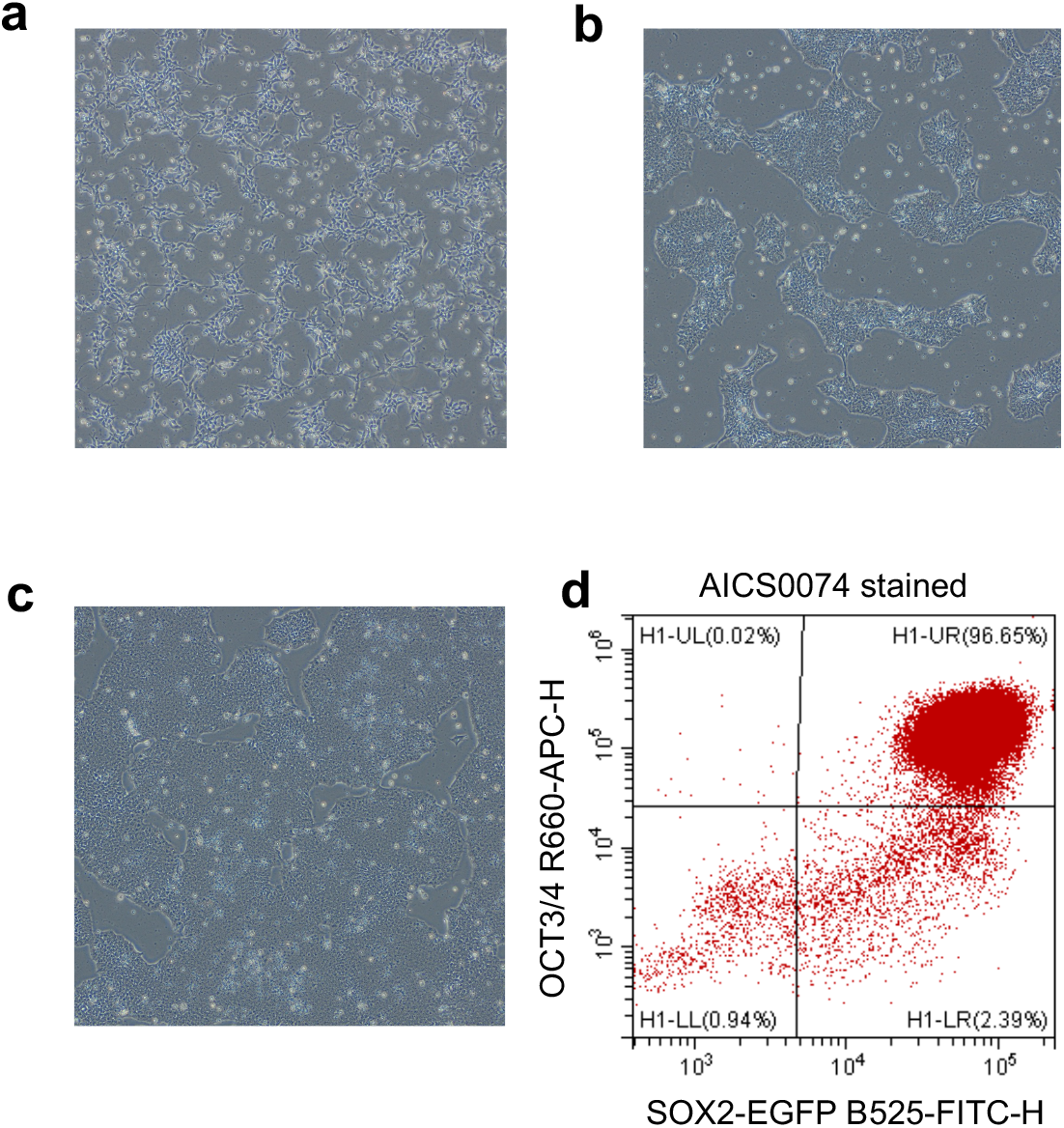
Routine maintenance and QC of hPSCs cell culture. **a**. hPSCs 1 day after seeding (In ROCK inhibitor). **b**. hPSCs 2 days after seeding. **c**. hPSCs 3 days after seeding, confluent enough to begin passaging. **d**. flow cytometry analysis of SOX2 and OCT3/4 expression shows 95%+ of cells in double positive state.

**Fig. S2.**
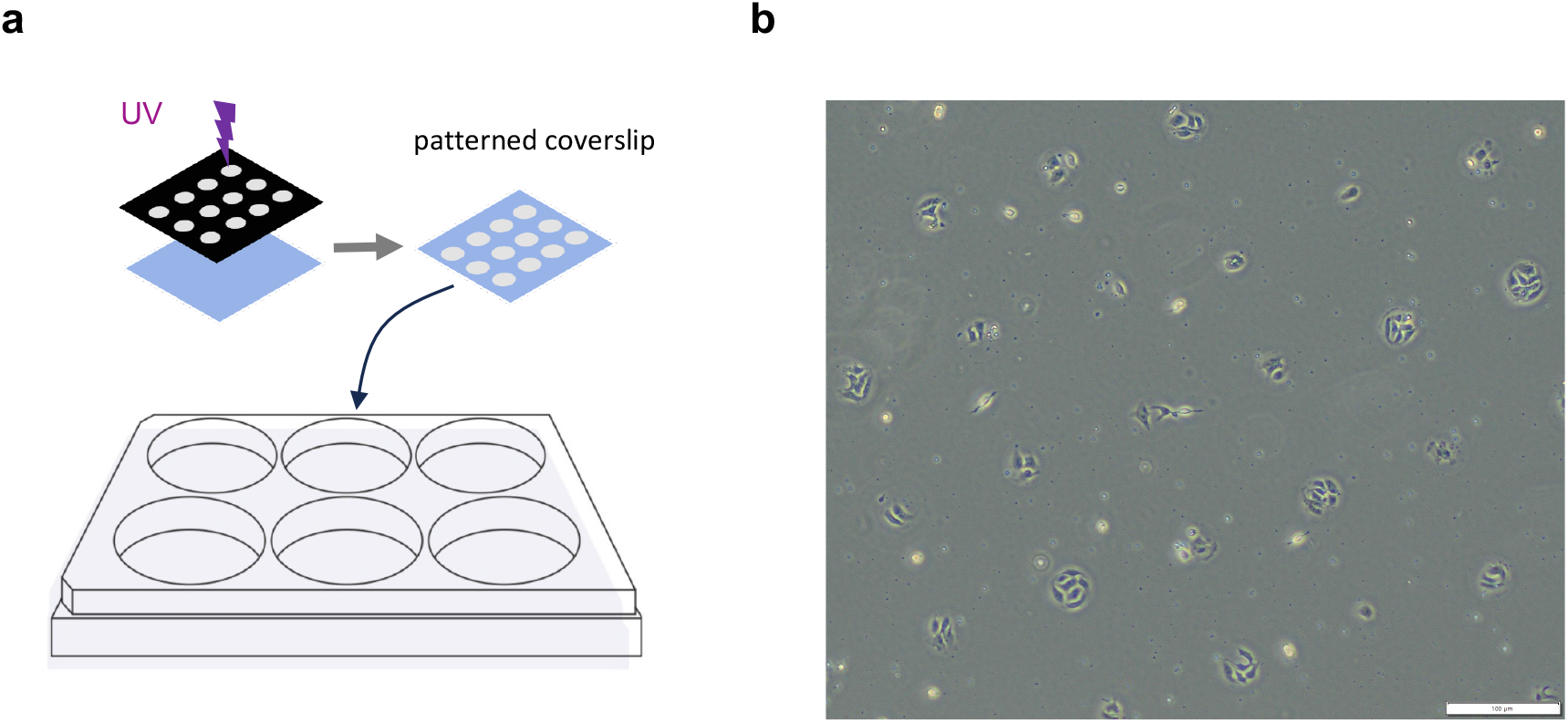
Preparation of adhesive islands and cell seeding. **a**. Patterned substrates were prepared on 22 × 22 mm glass coverslips and transferred to 6-well plates. **b**. Example of seeding cell on adhesive islands with the radius of R=25μm (seeding density of 50K cells per well). Scale bar shows 100 μm

**Fig. S3.**
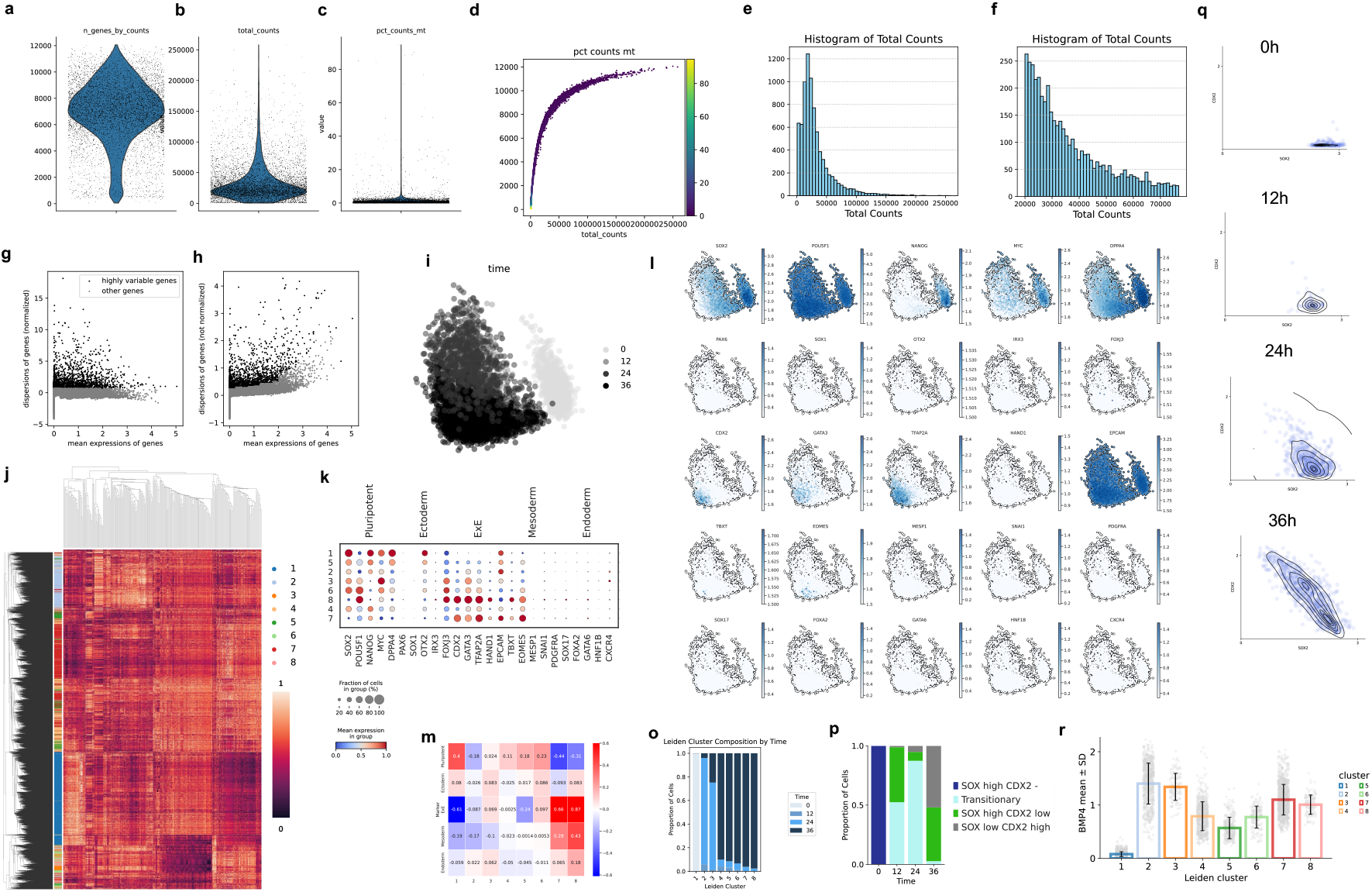
Single cell analysis of cell state transition. **a-c**. Violin plots showing quality control metrics per cell, including number of genes detected (a), total UMI counts (b), and percent mitochondrial transcripts (c). **d**. Joint scatter plot of mitochondrial content vs. total UMI counts with cells colored by % mitochondrial transcripts. **e-f**. Histograms of total counts before (e) and after (f) filtering to retain cells within the 20,000-77,000 UMI range. **g-h**, Mean vs dispersion plots of gene expression after (g) and before (h) normalization; black points indicate top 2,000 highly variable genes (HVGs) used for downstream analysis. **i**. PCA projection of all cells colored by timepoint (0, 12, 24, 36 hr after BMP4 stimulation), showing a temporal continuum. **j**. Cluster map of PCA neighborhood graph colored by Leiden cluster identity and expression of HVGs. Clustering was performed on the top 50 PCs using 50 neighbors and resolution = 0.6. **k**. Dot plot summarizing expression of key marker genes across clusters. Dot size reflects the proportion of cells expressing each gene, and color denotes mean expression. Markers cover pluripotency, ectoderm, mesoderm, endoderm, and extraembryonic fates. **l.** PCA overlays of gene expression for marker genes. The plots are visualized for imputed gene expression (MAGIC, Knn=2). **m**. Cluster-level summary of lineage marker group score. **o**. Stacked bar plot showing the composition of each Leiden cluster across timepoints. **p**. Stacked bar plot showing the composition of annotated groups across timepoints. **q**. Kernel density estimate (KDE) plots of SOX2 and CDX2 imputed expression (MAGIC, Knn=2) at 0, 12, 24, and 36 hr post-BMP4. Each point represents a cell, and contours indicate local cell density, revealing progressive bifurcation along the fate axis. **r**. Bar plot of BMP4 expression (log transformed) in each cluster.

**Fig. S4.**
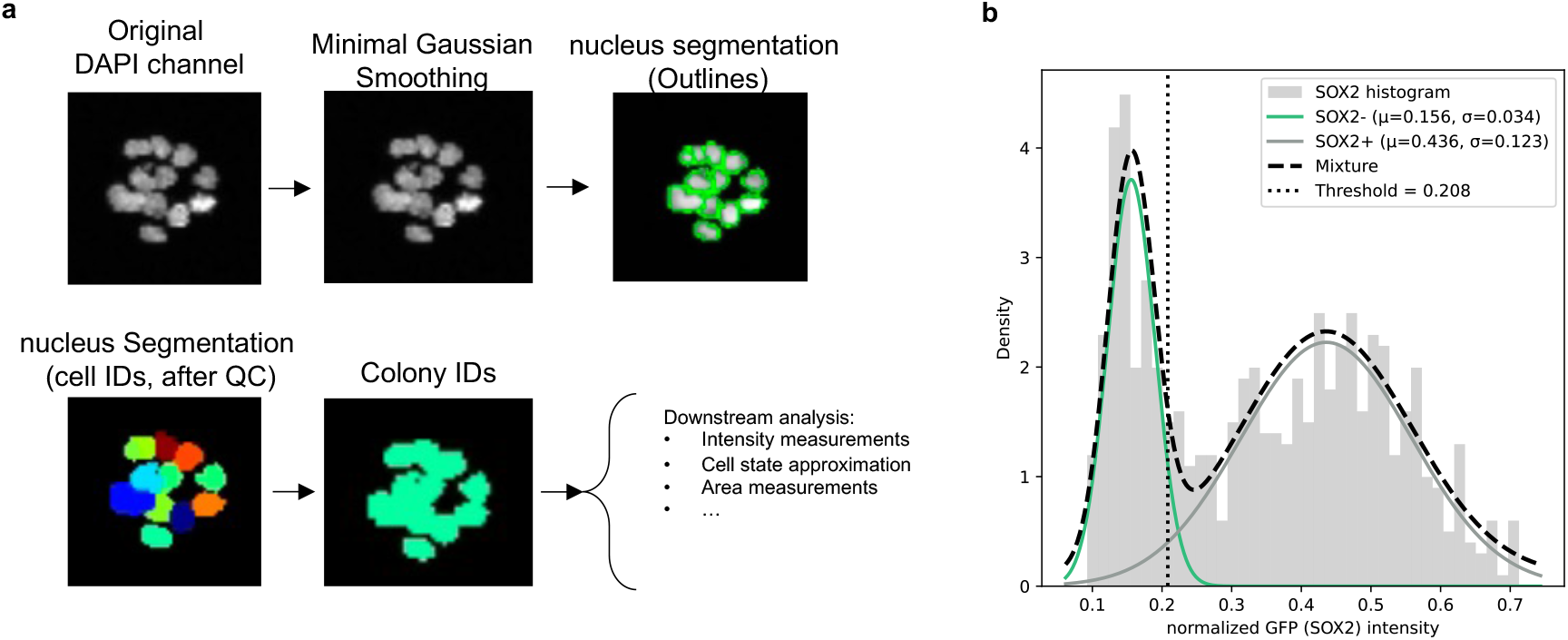
Cell segmentation and cell state binarization. **a**. Representative image processing workflow: DAPI channel is smoothed and segmented to identify individual nuclei and assign cell and colony IDs. **b**. Measurement of SOX2-GFP intensity (normalized). Plot shows fitted Gaussian mixture separating SOX2- and SOX2+ populations. k-means clustering threshold (dotted line) was used to determine SOX2 expression state.

**Fig. S5.**
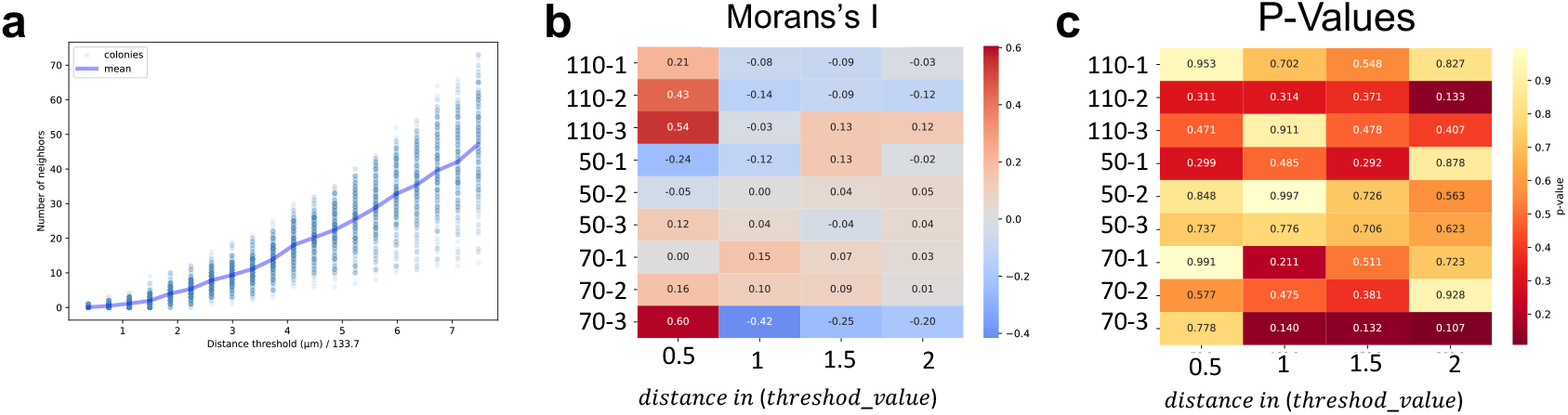
Inter-colony spatial coordination in SOX2 expression assessed via Moran’s I. **a**. Representative example of relationship between distance threshold and number of neighboring colonies used to construct spatial graphs. Each dot represents the number of neighbors detected for one colony at the given threshold, with the blue line showing the mean across colonies. **b**. Moran’s I statistic computed for each experimental replicate at multiple spatial thresholds (thresholds calculated per sample). Warmer colors indicate stronger positive spatial autocorrelation (coordinated SOX2 expression across neighboring colonies); cooler values indicate negative or anti-correlated patterns. **c**. Corresponding p-values obtained from permutation testing (999 shuffles per replicate and threshold). No consistent significance was observed across replicates, suggesting weak or heterogeneous spatial coordination at the inter-colony level.

**Fig. S6.**
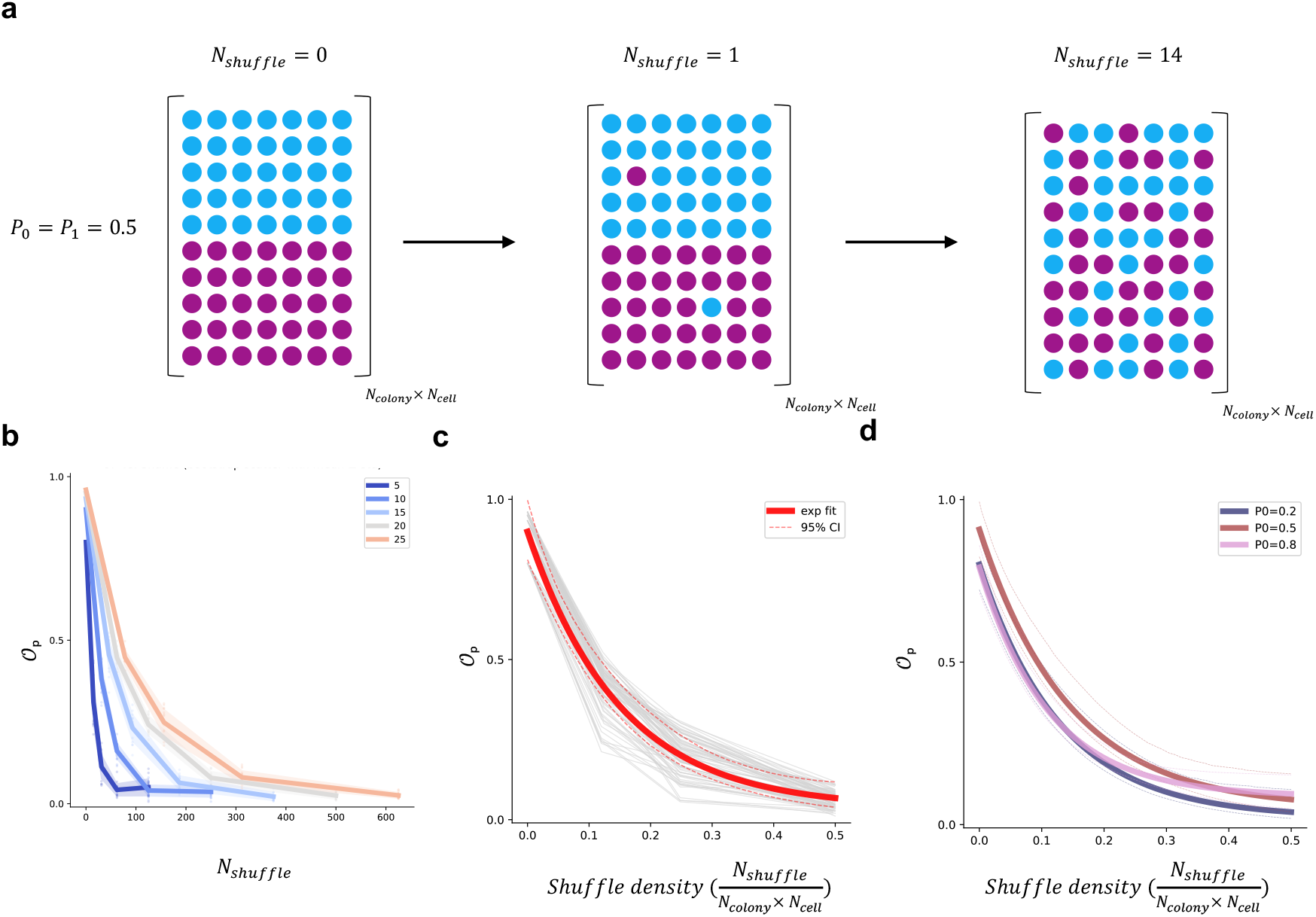
Characterization of 𝒪_***p***_ via disorder-induced permutations. **a**. Schematic of colony ensembles with increasing cross-colony shuffle steps. Each colony contains *N*_*cell*_ binary-state cells, initialized in an ordered configuration (as uniform-state as possible). Cross-colony swaps are only accepted between cells of differing states, preserving global distribution while increasing disorder. **b**. Decay of the order parameter 𝒪_*p*_ with increasing colony size and shuffle count. Points represent mean ± s.d. across bootstraps. **c**. When plotted against normalized shuffle density *N*_*shuffle*_/(*N*_*colony*_ × *N*_*cell*_), all curves collapse onto a single master curve. Red line shows an exponential fit with 95% confidence interval. **d**. Collapse persists across different global state fractions *P*_0_ ∈ {0.2, 0.5, 0.8}, demonstrating robustness of *𝒪*_*p*_ as a scale-invariant measure of multicellular order.

**Fig. S7.**
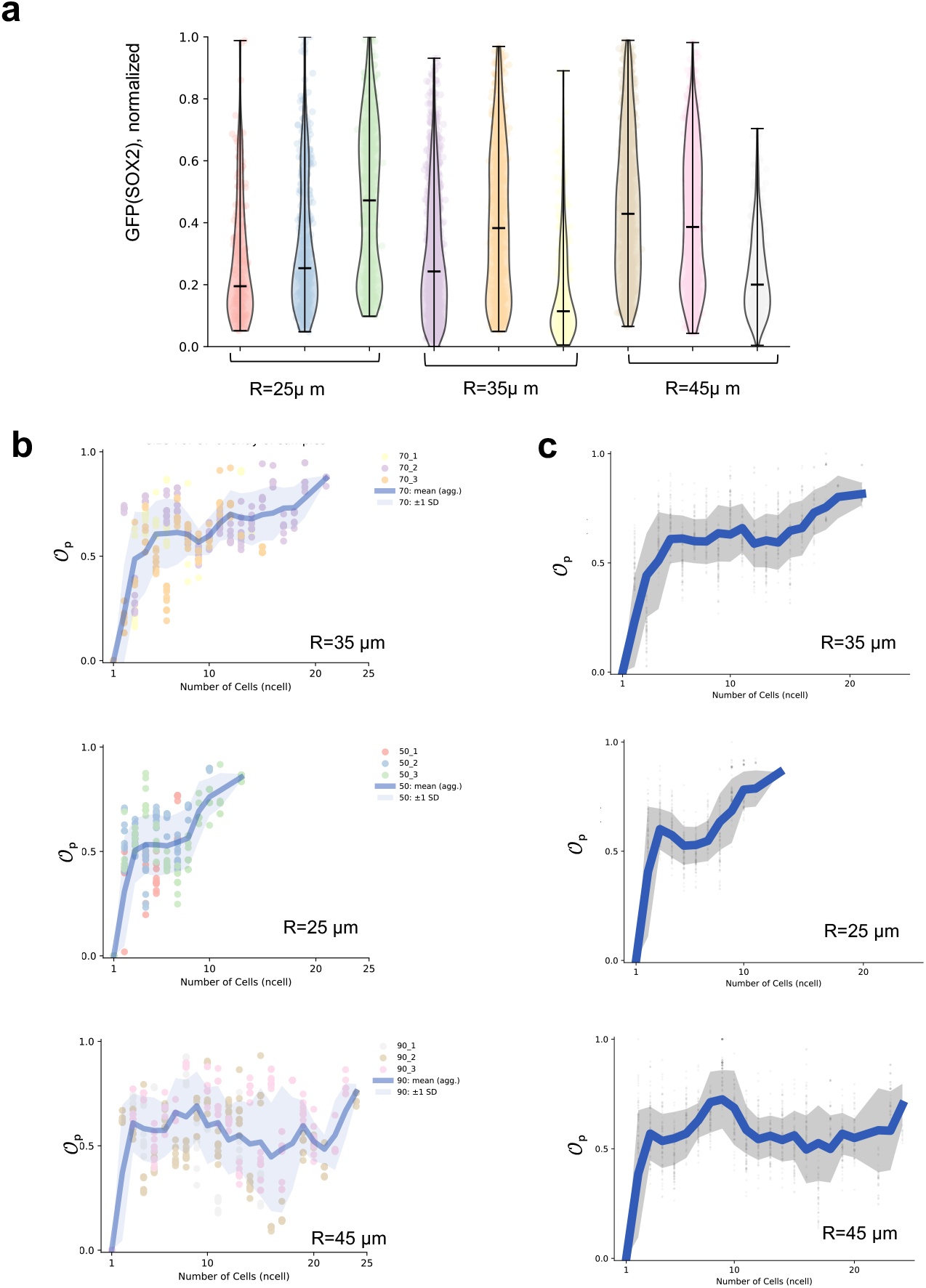
Estimation of order parameter *𝒪*_*p*_ across colony sizes and experimental replicates. **a**. Distribution of normalized SOX2-GFP intensities across colony sizes and experimental replicates for each pattern diameter (R = 25 µm, 35 µm, 45 µm). Each violin shows the distribution across colonies of a given size in one replicate (separate chip with multiple colonies). **b**. Order parameter *𝒪*_*p*_ curves for each radius condition. Dots show individual bootstrap samples (N = 10), each computed on 60% of colonies sampled with replacement from each replicate (colored). Solid line and shaded region show the mean ± s.d. across bootstraps. **c**. Final pooled *𝒪*_*p*_ curves combining all 3 experimental replicates per condition. A sliding window of width 3 was applied across colony sizes to compute smoothed means and standard deviations. For single-cell colonies, *𝒪*_*p*_ = 0 by definition, as coordination is not meaningful. Dots show individual bootstrap samples (N = 100).

**Fig. S8.**
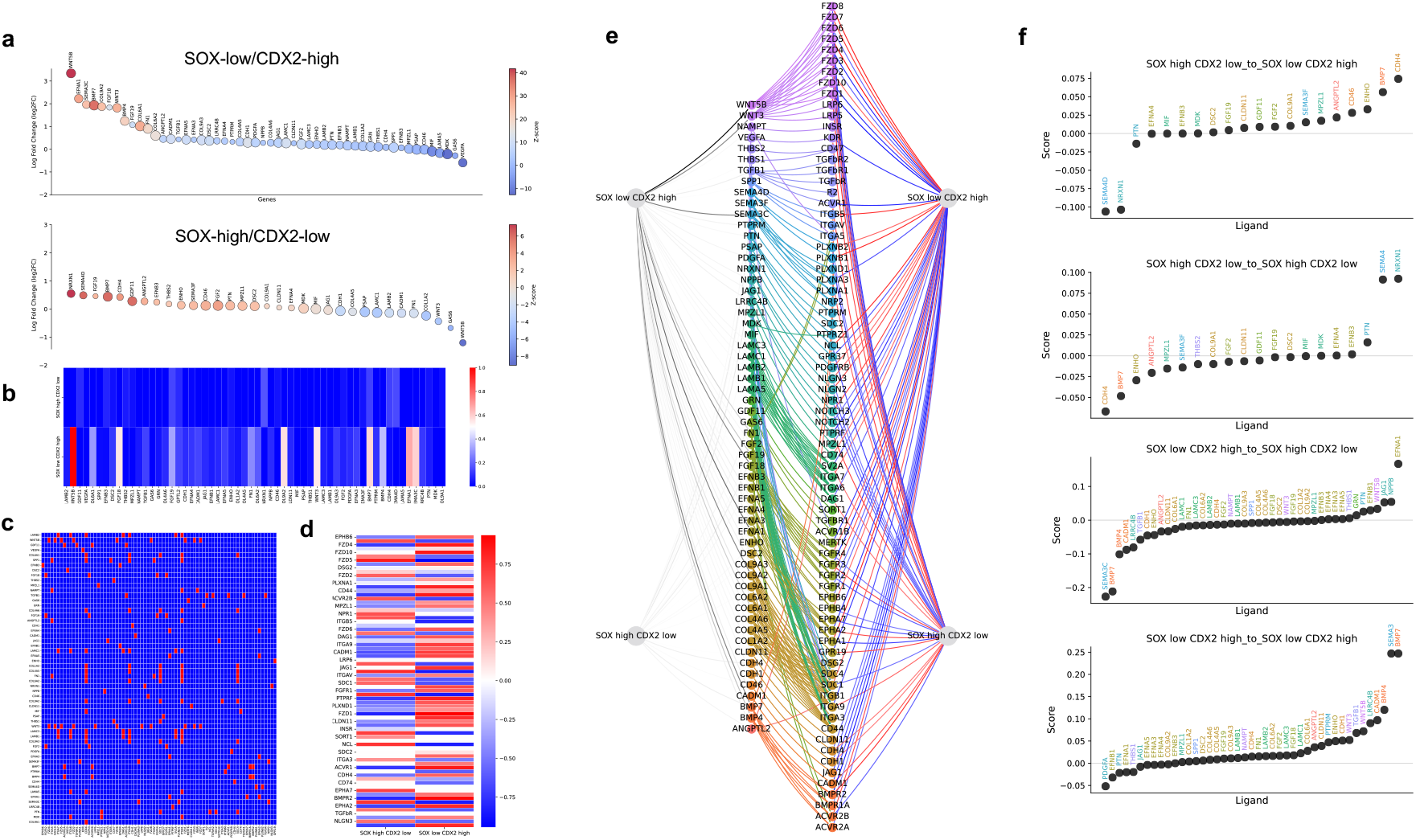
Inference of ligand-receptor-TF signaling paths between transcriptionally distinct cell states. **a**. Top ligands differentially expressed in “SOX-low/CDX2-high” (top) and “SOX-high/CDX2-low” (bottom) source populations, computed using log2 fold-change and used to construct the ligand usage matrix *M*_1_. **b**. Normalized ligand usage matrix *M*_1_, where each row is a source cluster and columns are ligands. Values range from 0 to 1, representing expression enrichment. **c**. Binary ligand-receptor adjacency matrix *M*_2_, indicating functional interactions derived from curated databases (CellChat + CellPhoneDB). **d**. Recepto-TF correlation matrix *M*_3_, combining DREMI and Pearson correlation of minimally imputed gene expression (MAGIC, knn=2). Only receptor-TF pairs with |Corr| > 0.5 are shown. **e**. Multi-layer signaling graph linking source clusters → ligands → receptors → target clusters. Edge colors denote signaling pathway families (e.g., WNT, BMP), and receptor→TF correlations are shown in blue-red scale. **f**. Ranked ligand scores for directional cluster pairs, summarizing top inferred communication paths (non-zero PathScore(*s, l, r, t*), (ligands are colored by pathway family).

**Fig. S9.**
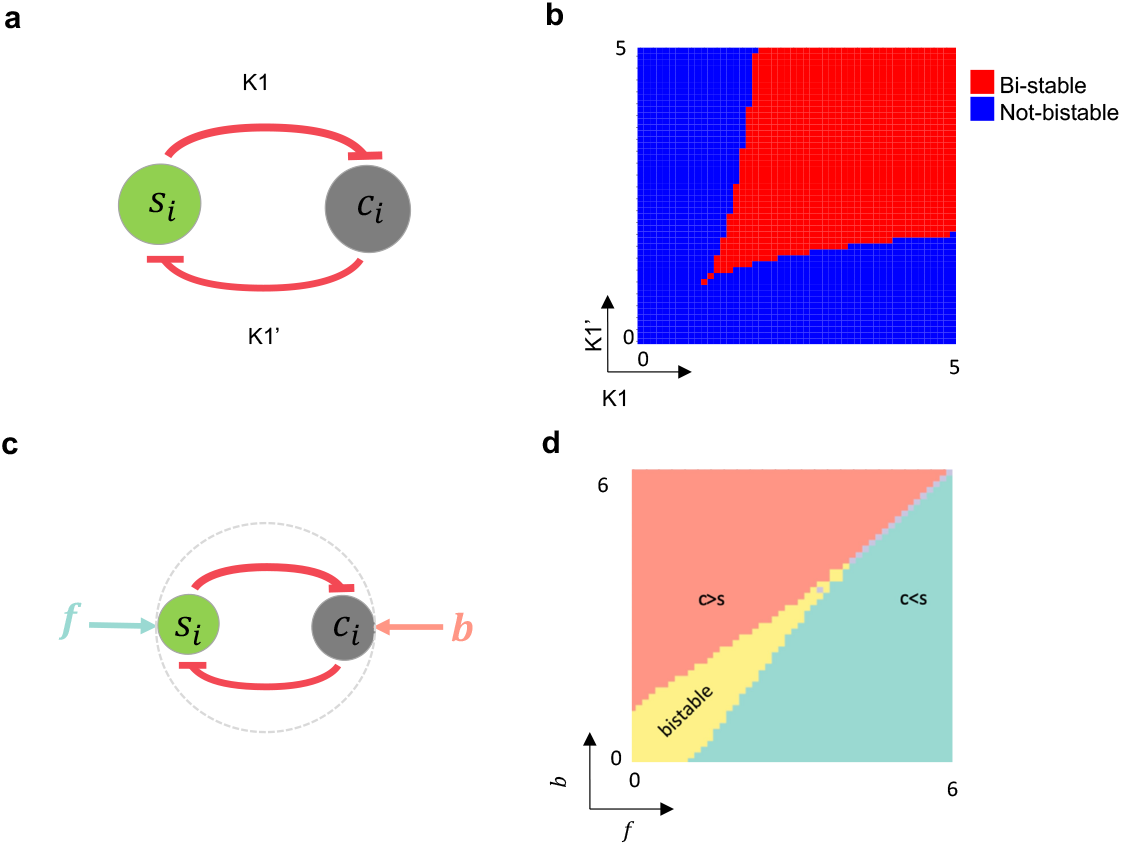
Single-cell Bistable switch and signaling extension. **a**. Schematic of mutual repression toggle between *s* and *c*. **b**. Parameter scan showing bistable (red) and monostable (blue) regimes as a function of production rates *K*1 and *K*1′. **c**. Extension of the toggle with constant level external factors *f* (promoting *s*) and *b* (promoting *c*). **d**. Phase diagram of system behavior as a function of the constant levels of (*f, b*) highlighting bistable (the toggle state remains unchanged) and biased regimes (external factors can change the state of the toggle switch).

**Fig. S10.**
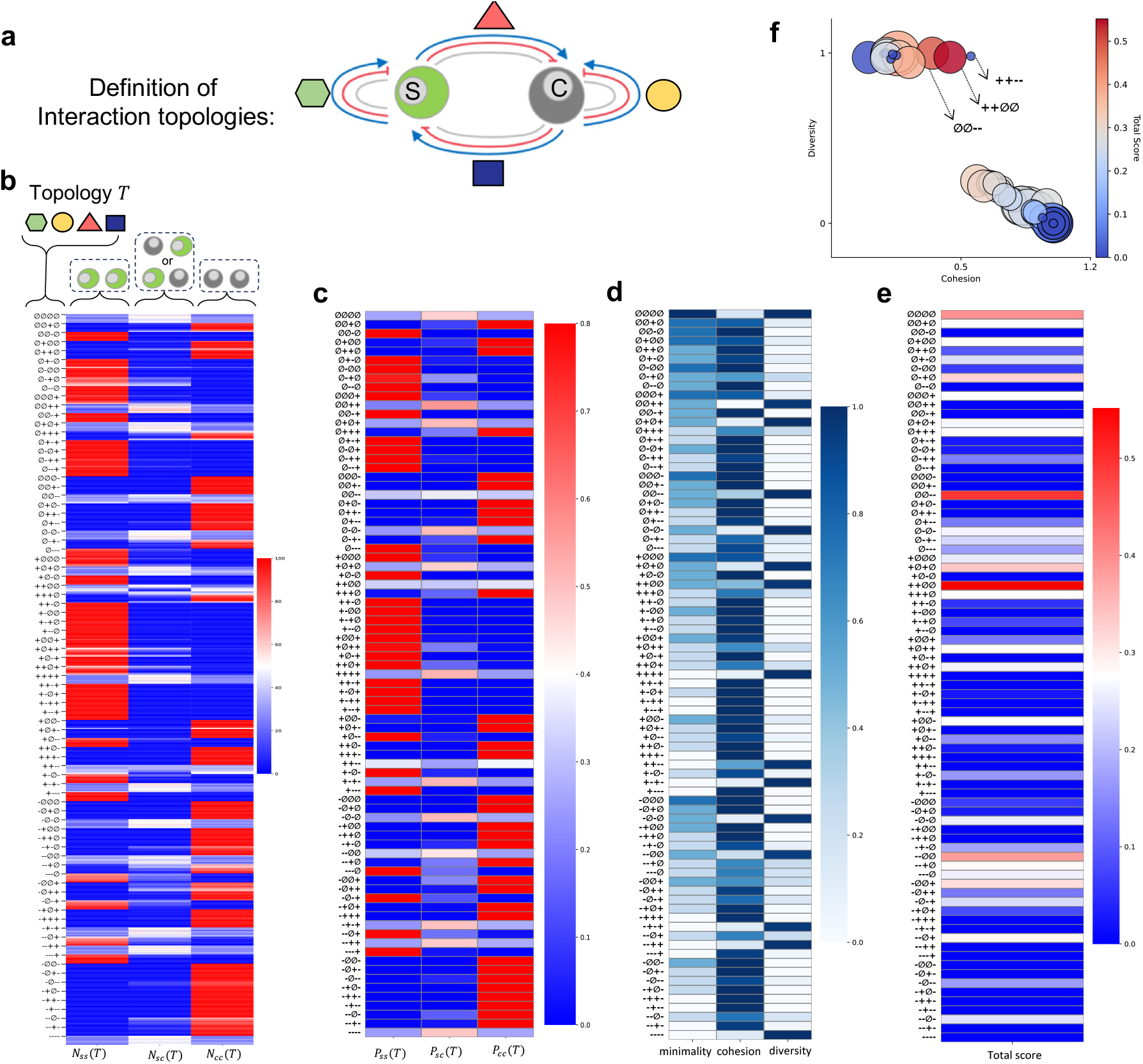
Scoring signaling topologies by minimality, cohesion, and fate balance. **a**. Schematic definition of two-cell signaling topologies, varying in interaction structure. Shapes represent the interactions and are referenced in panel (b). Each interaction can be positive (+), negative (-) or empty (∅). **b**. Counted outcomes for each topology *T* based on 100 random initializations. and 8 different values of 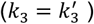 in the range of [1,2]. Each block of the row shows the empirical probability of reaching SS (SOX2-SOX2), CC (CDX2-CDX2), or SC (mixed) fates. **c**. Pooled out comes of (b). Results are aggregated and averaged and converted to probabilities for all range of parameters. **d**. Normalized metrics computed from these probabilities: *Minimality* favors sparse topologies, *Cohesion* measures agreement between cells, and *Diversity* quantifies balance between fates. **e**. Composite score combining all metrics by geometric mean, enabling ranking of topologies that are parsimonious, stable, and fate-balanced. **f**. Evaluation of all topologies across three axes: diversity (y-axis), cohesion (x-axis), and minimality (circle size, inverse of interaction count). Color reflects overall score (red = high, blue = low). A small subset of topologies is illustrated by arrow.

**Fig. S11.**
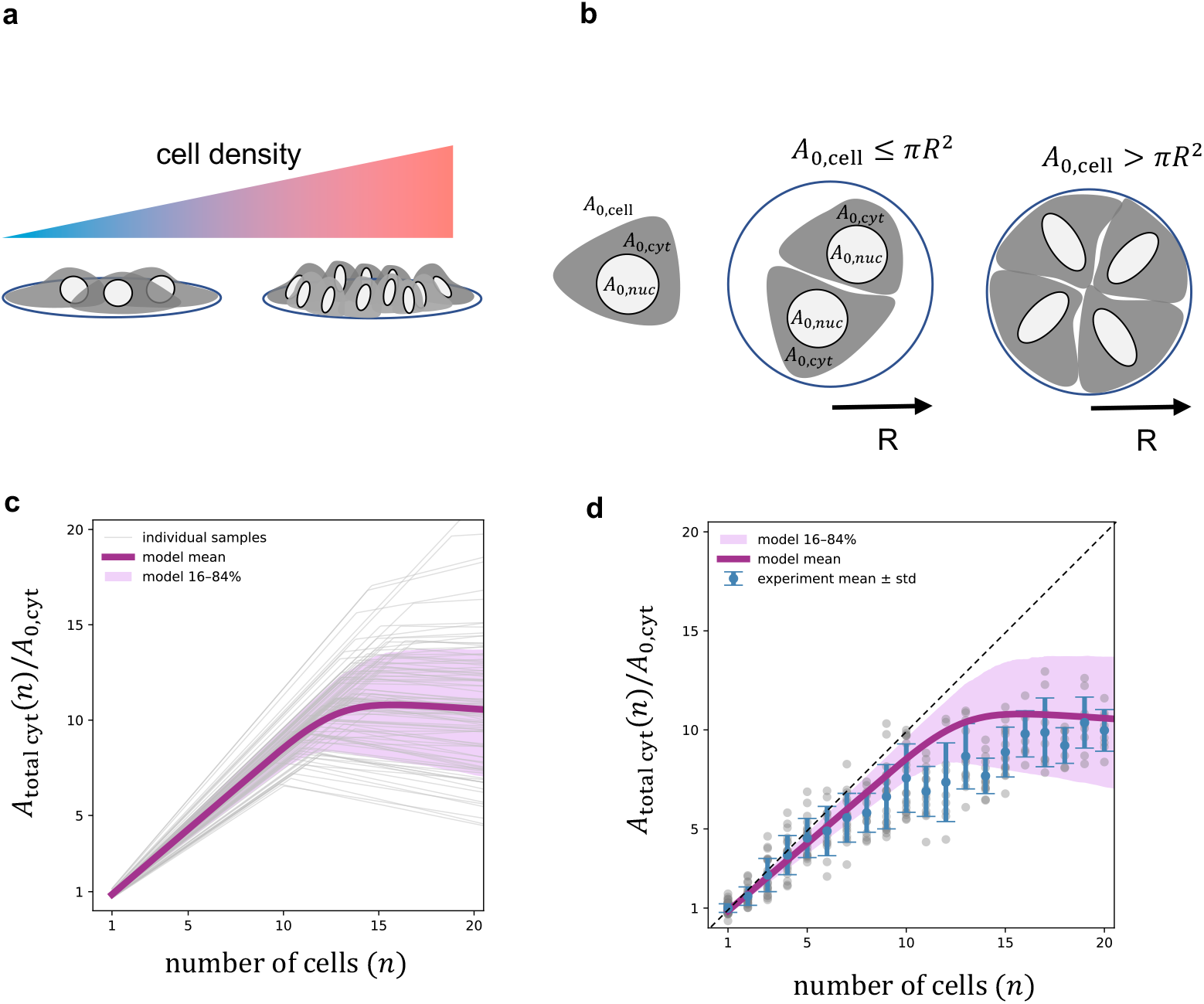
Mechanical packing model of extra-nuclear reservoir. **a**. Schematic of increasing colony density and confinement. **b**. Two regimes of cytoplasmic area depending on available pattern area **c**. Model predictions of normalized total cytoplasmic area as a function of cell number, showing linear and confined regimes. **d**. Comparison of model predictions with experimental measurements (mean ± s.d.), (χ^2^ = 0.54).

**Fig. S12.**
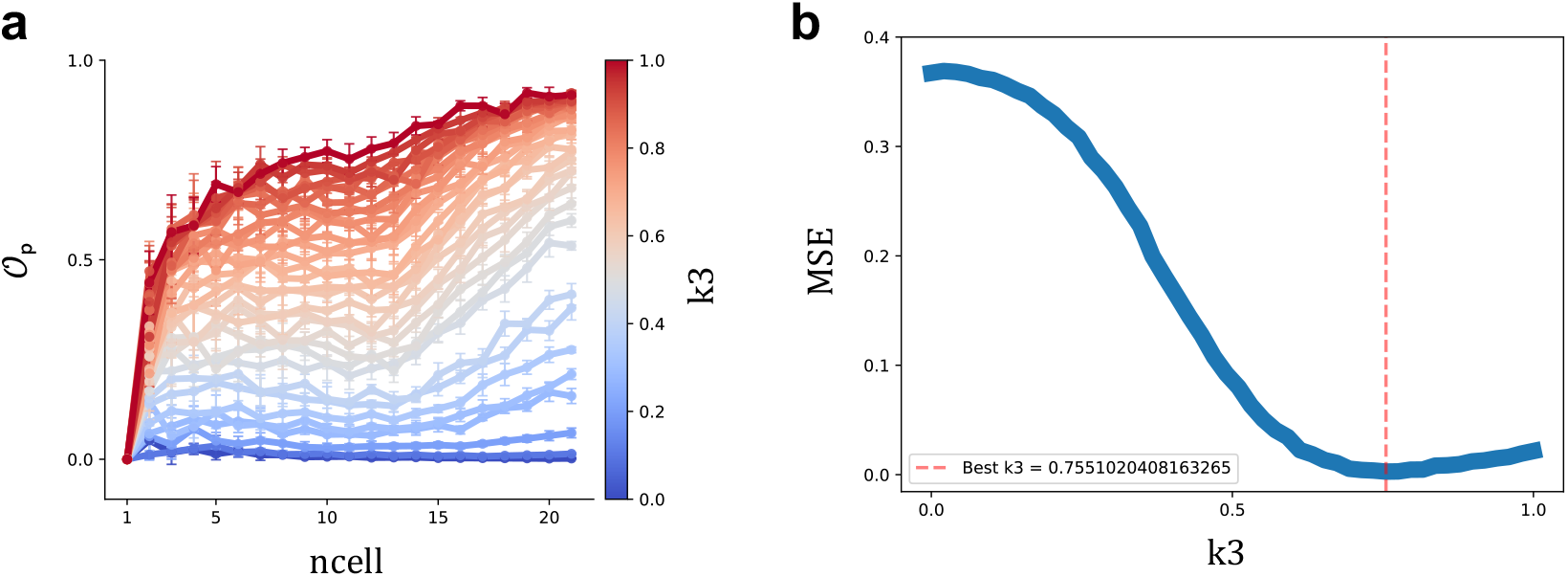
Fitting the free parameter k3 using order-parameter data. a. Simulation sweep of k3 (color bar) showing the predicted order parameter (*𝒪*_p_) versus colony size (number of cells in the colony). **b**. Model-vs-experiment mean-squared error (MSE) computed between the experimental *𝒪*_p_ means and each simulated mean curve across size bins. The minimum defines the best-fit parameter k3≈0.75 (shown by vertical dashed line). All other parameters were fixed from independent measurements.

**Fig. S13.**
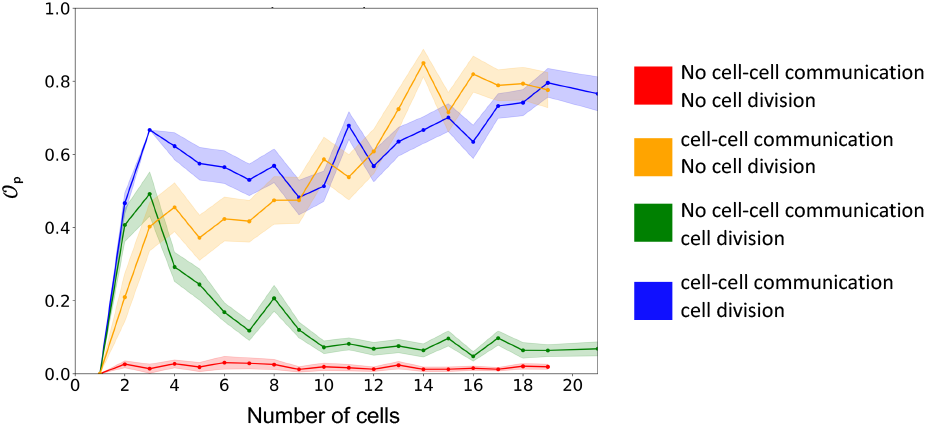
Effect of cell division and cell-cell communication on colony-level order. Order parameter *𝒪*_*p*_ is plotted against colony size under four simulation regimes: (red) no communication or division; (orange) communication only; (green) division only; (blue) both communication and division. Cell-cell communication alone substantially increases order. Cell division alone slightly increased order in small colonies by amplifying early stochastic differences.

**Fig. S14.**
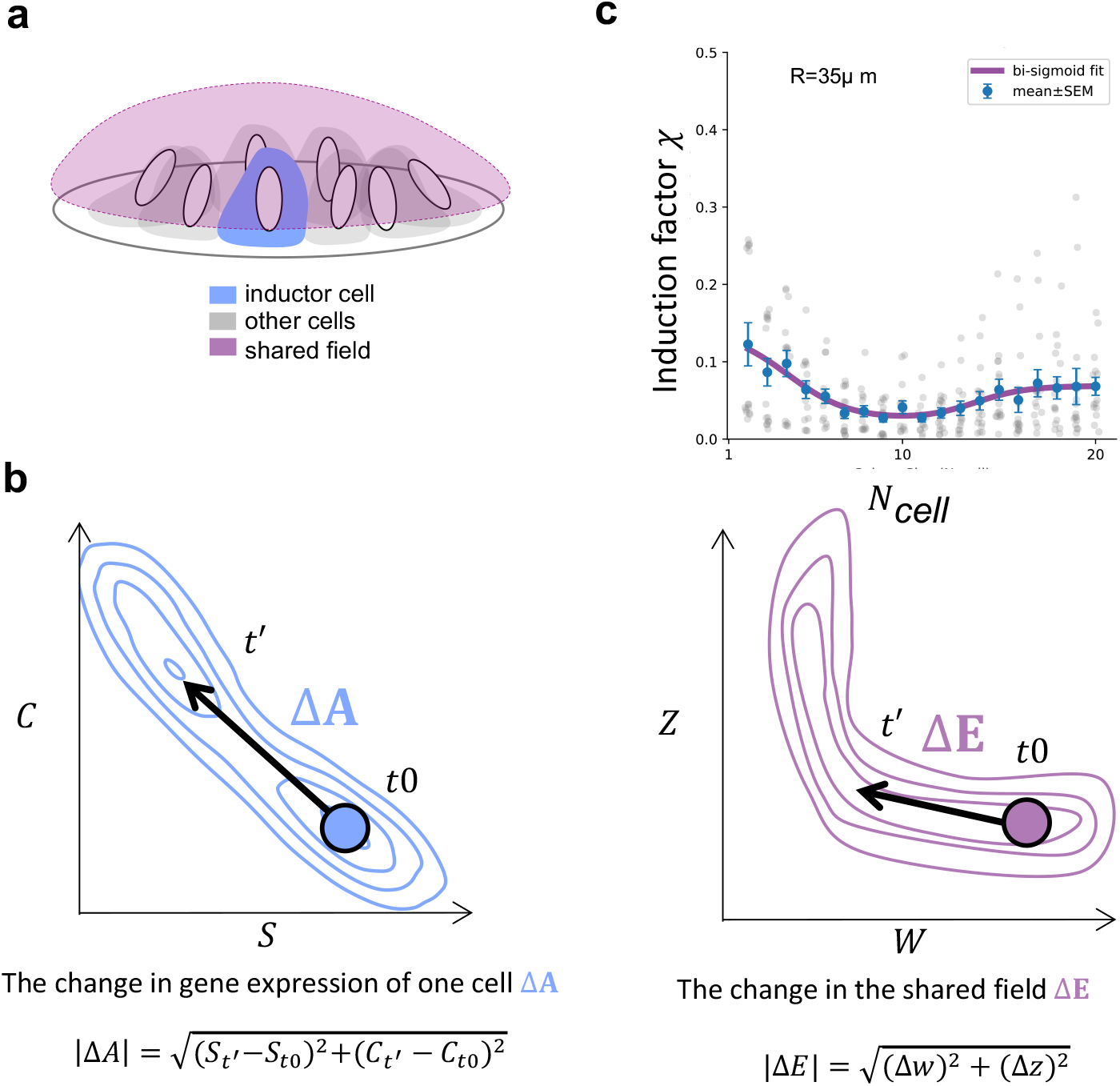
Quantifying cross colony information propagation via induction factor. **a**. Schematic of a perturbed inductor cell (blue) altering the shared signaling field (purple) within a colony. **b**. Geometric definitions of the inductor’s change in gene expression Δ*A* and the corresponding shift in the shared field Δ*E*, computed as Euclidean displacements between time-averaged pre-pulse and post-pulse states. The induction factor χ =∣ Δ*E* ∣/∣ Δ*A* ∣ captures how strongly a local perturbation propagates into the collective environment. **c**. Induction factor as a function of colony size (*N*_*cell*_) for different colony radii. Each dot is a simulation replicate; solid lines show bi-sigmoid fits capturing the non-monotonic behavior of χ(*N*).

**Fig. S15.**
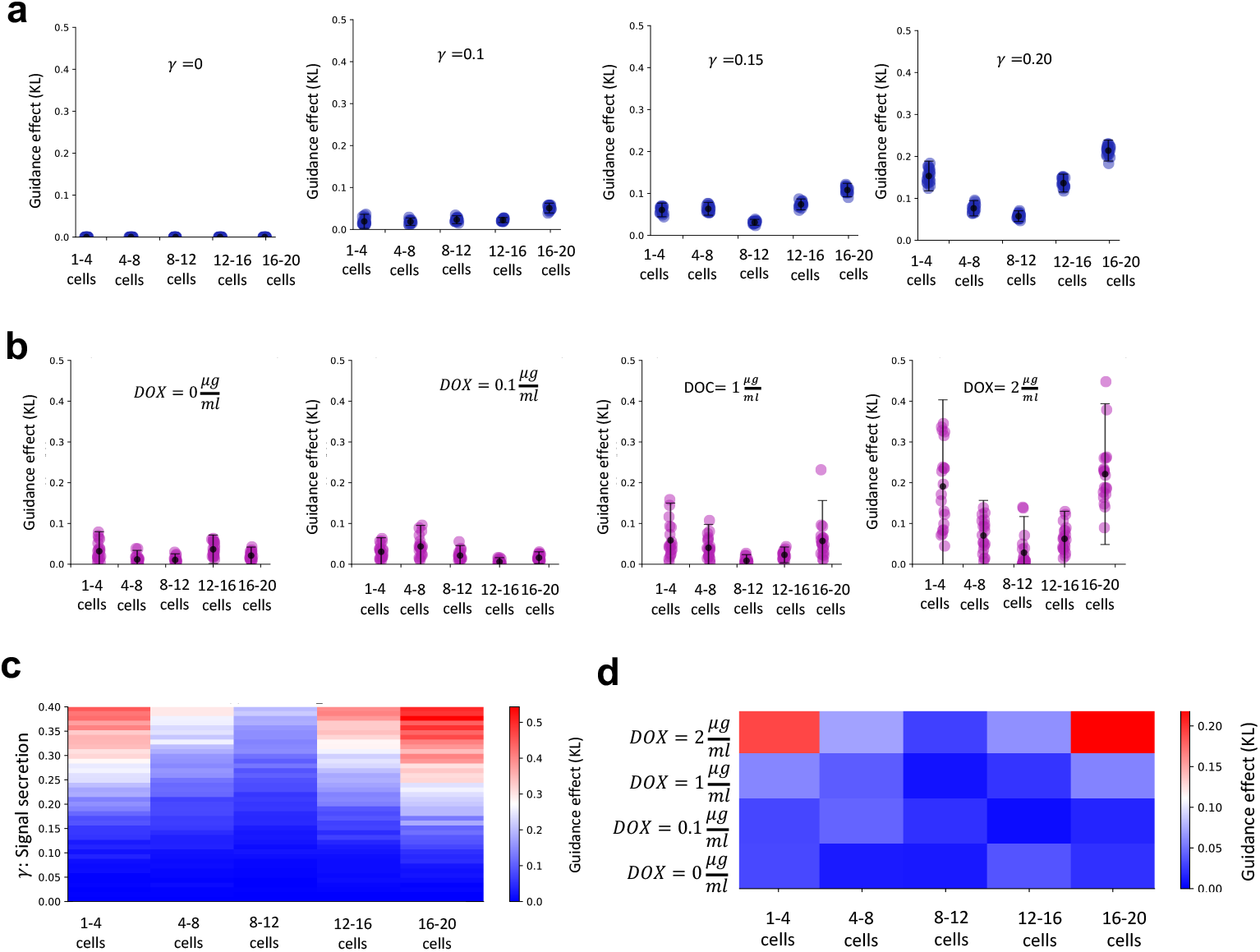
Guidance effect across signal strength and colony size: model vs experiment. **a**. Simulations with one guide cell: guidance effect is quantified as KL divergence between SOX2 distributions with vs without a guide cell, within each colony-size bin. Each dot is a bootstrap sample; vertical bars show mean ± SD. Increasing γ strengthens guidance and reveals a biphasic size dependence. **b**. DOX-inducible BMP4 guide-cell: each dot is a bootstrap sample of KL (with vs without one guide cell) for the indicated DOX dose; bars are mean ± SD. Guidance emerges at higher DOX inductions and follows the same biphasic size profile as model predicted. **c**. Model summary heatmap of mean KL over bootstraps versus colony size (x-axis) and γ (y-axis), highlighting susceptibility windows at small and large sizes. **d**. Experimental summary heatmap of mean KL over bootstraps versus colony size and DOX dose, confirming strongest effects at 2 μg mL^−1^ in the smallest and largest colonies.

**Fig. S16.**
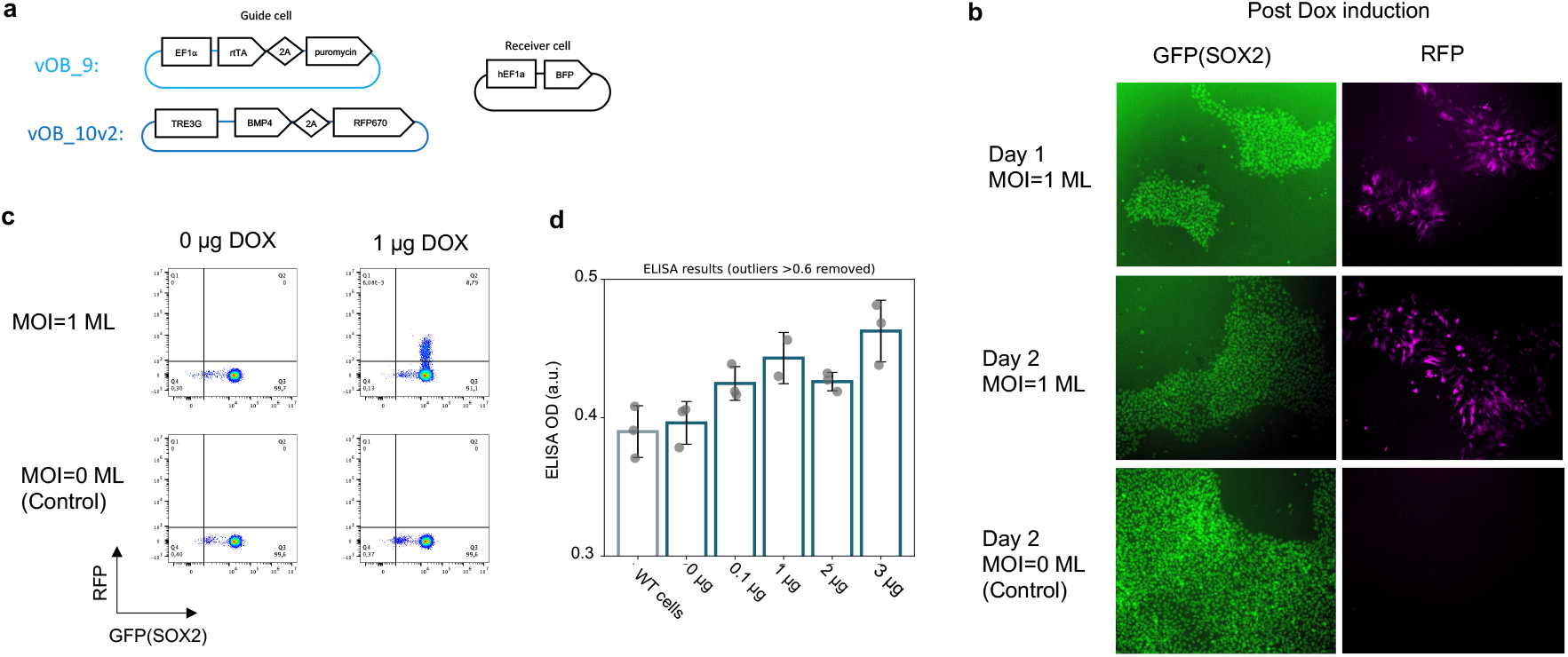
Preparation of guide cell experiment. **a**. Schematic of lentiviral constructs for guide cells (vOB_9, vOB_10v2) and receiver cells. **b**. Fluorescence microscopy showing DOX-inducible RFP expression in guide cells (MOI=1) alongside GFP(SOX2) at Day 1-2 post DOX induction. **c**. Flow cytometry validation of DOX activation of RFP induction in guide cells, Day 2 post induction (MOI=1 vs. MOI=0 control). **d**. Absorbance from BMP reporter assay (ELISA) increases with DOX induction. One data point of technical readout (belonging to DOX=1μg) has been removed (outlier value= 0.65).

**Fig. S17.**
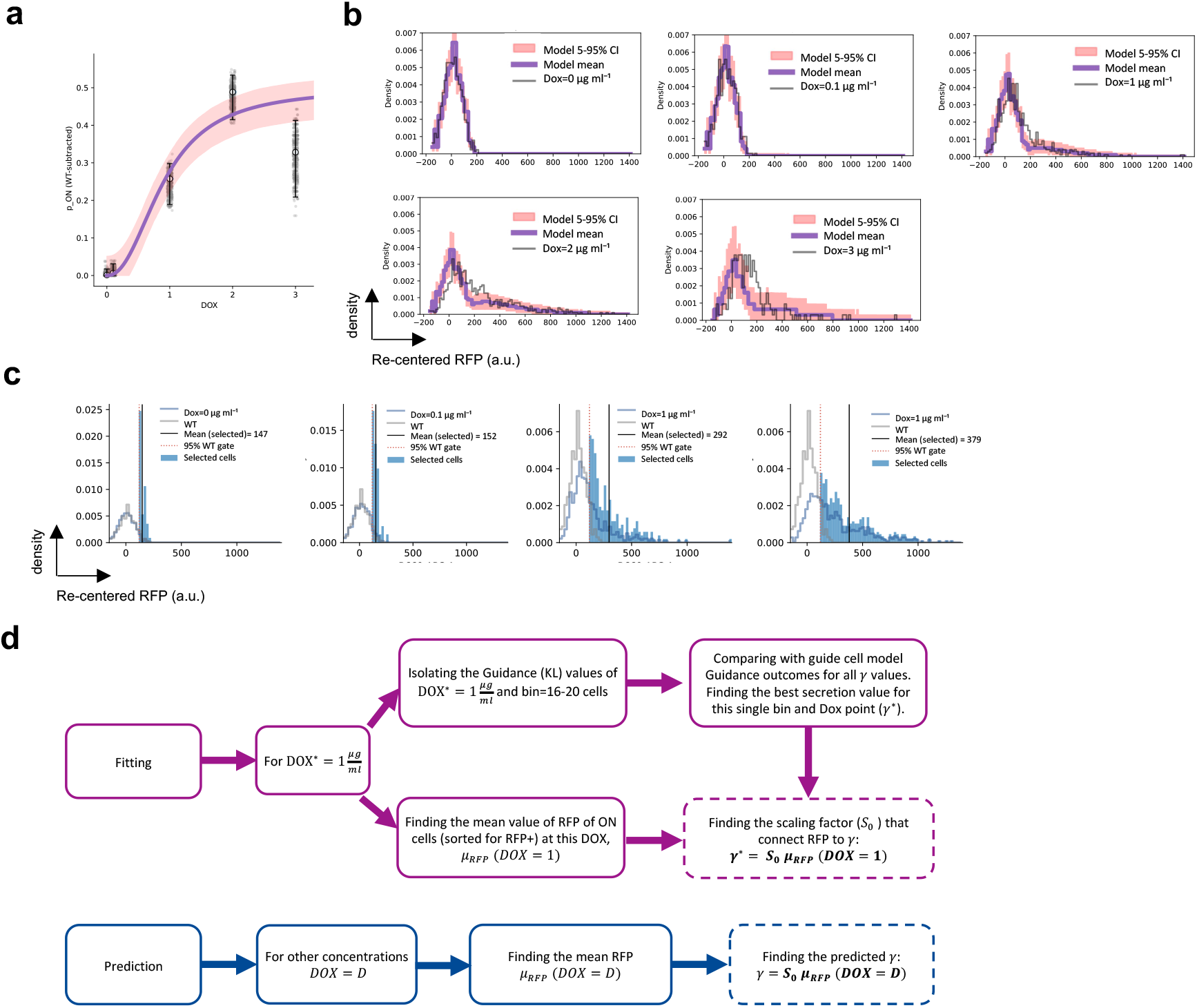
Connecting DOX-inducible BMP4-2A-RFP measurements to the guide-cell model. **a**. Fraction of ON cells vs DOX. ON is defined by a fixed RED-channel gate set at the 95th percentile of WT (no-transduction) cells. A simple Poisson-Hill activation model was fit only to these ON-fraction data; the curve shows the fit and the band shows the 95% CI. Best-fit parameters: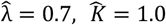, and 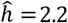. **b**. Full RFP distributions across DOX doses. Grey histograms are experiments; overlaid lines and shaded bands are predictions from the model in a. The model captures the dose-dependent shift and broadening of RFP. All RFP values have been shifted to center WT (background). **c**. Gated ON populations used to compute the mean RFP at each dose. The vertical dotted line marks the WT-derived 95% gate; shaded bars indicate selected cells. Reported means are used as the experimental proxy for secretion. The RFP values have been shifted to center WT (background). **d**. Calibration of secretion in the guide-cell model. Using a single anchor condition (DOX = 1 μg mL^−1^, 16-20-cell bin), we chose the secretion factor that best matched the experimental guidance (KL) at that condition (*γ*^∗^ =1.4). This defines a proportional mapping from the mean RFP at any dose to the model’s secretion factor, enabling direct mode-experiment comparisons of guidance across DOX.

